# Kaposi’s sarcoma-associated herpesvirus ORF67.5 functions as a component of the terminase complex

**DOI:** 10.1101/2023.03.29.534847

**Authors:** Yuki Iwaisako, Tadashi Watanabe, Youichi Suzuki, Takashi Nakano, Masahiro Fujimuro

## Abstract

Kaposi’s sarcoma-associated herpesvirus (KSHV) is a double-stranded DNA gammaherpesvirus, with a poorly characterized lytic replication cycle. However, the lytic replication cycle of the alpha- and betaherpesviruses are well characterized. During lytic infection of alpha- and betaherpesviruses, the viral genome is replicated as a precursor form, which contains tandem genomes linked via terminal repeats (TRs). One genomic unit of the precursor form is packaged into a capsid and is cleaved at the TR by the terminase complex. While the alpha- and betaherpesvirus terminases are well characterized, the KSHV terminase remains poorly understood. KSHV ORF7, ORF29, and ORF67.5 are presumed to be components of the terminase complex based on their homology to other terminase proteins. We previously reported that ORF7-deficient KSHV formed numerous immature soccer ball-like capsids and failed to cleave the TRs. ORF7 interacted with ORF29 and ORF67.5, however ORF29 and ORF67.5 did not interact with each other. While these results suggested that ORF7 is important for KSHV terminase function and capsid formation, the function of ORF67.5 was completely unknown. Therefore, to analyze the function of ORF67.5, we constructed ORF67.5-deficient BAC16. ORF67.5-deficient KSHV failed to produce infectious virus, failed to cleave the TRs, and numerous soccer ball-like capsids were observed in ORF67.5-deficient KSHV-harboring cells. Furthermore, ORF67.5 promoted the interaction between ORF7 and ORF29, and ORF29 increased the interaction between ORF67.5 and ORF7. Thus, our data indicated that ORF67.5 functions as a component of the KSHV terminase complex by contributing to TR cleavage, terminase complex formation, capsid formation, and virus production.

**IMPORTANCE:** Although the formation and function of the alpha- and betaherpesvirus terminase complexes are well understood, the KSHV terminase complex is still largely uncharacterized. This complex presumably contains KSHV ORF7, ORF29, and ORF67.5. We were the first to report the presence of soccer ball-like capsids in ORF7-deficient KSHV-harboring lytic-induced cells. Here, we demonstrated that ORF67.5-deficient KSHV also formed the soccer ball-like capsids in lytic-induced cells. Moreover, ORF67.5 was required for TR cleavage, infectious virus production, and enhancement of the interaction between ORF7 and ORF29. ORF67.5 has several highly conserved regions among its human herpesviral homologs. These regions were necessary for virus production and for the interaction of ORF67.5 with ORF7, which was supported by the AI-predicted structure model. Importantly, our results provide the first evidence showing that ORF67.5 is essential for terminase complex formation and TR cleavage.

## Introduction

KSHV is a double-stranded DNA human herpesvirus belonging to the gammaherpesvirinae subfamily (1). Other human herpesviruses including KSHV are categorized into three subfamilies: alphaherpesvirinae [herpes simplex virus 1 (HSV-1), herpes simplex virus 2 (HSV-2), and varicella-zoster virus (VZV)], betaherpesvirinae [human cytomegalovirus (HCMV), human herpesvirus 6 (HHV-6), and human herpesvirus 7 (HHV-7)], and gammaherpesvirinae [Epstein-Barr virus (EBV) and KSHV] (2). KSHV is the etiologic agent of Kaposi’s sarcoma, primary effusion lymphoma, multicentric Castleman’s disease, and KSHV-associated inflammatory cytokine syndrome (3, 4, 5, 6). KSHV has two states of infection, latent and lytic. The expression of a viral protein referred to as the replication and transcription activator (RTA/ORF50), induces the transition from latent to lytic infection, which produces progeny virions (7). The KSHV genes that are expressed during lytic infection are classified as immediate early (IE), delayed early (DE), and late (L), based on differences in the timing of expression and the conditions required for expression (2). During lytic infection, viral capsids are formed in the nucleus of the host cell and the replicated viral genome is packaged into capsids to form a mature capsid. The mechanism responsible for mature capsid formation is still largely unknown for KSHV, but it is relatively well understood for HSV-1.

HSV-1 forms procapsids, A-capsids, B-capsids, and C-capsids during mature capsid formation (8, 9). First, a porous, hollow procapsid is formed with a globular outer layer consisting of major capsid proteins and a globular inner layer consisting of scaffold proteins (10, 11). The outer layer structure of the procapsid is supported by scaffold proteins, and the outer layer and scaffold proteins are linked together (12). Subsequently, when the link between the outer layer of the procapsid and the scaffold proteins is detached by the viral protease VP24, a globular scaffold inner layer remains inside the outer layer, and the outer layer undergoes spontaneous angularization to form a regular icosahedron. This regular icosahedron capsid, harboring a globular inner layer, is referred to as the B-capsid (13, 14, 15). Since both procapsids and B-capsids harbor a globular scaffold protein layer, it is difficult to distinguish between them in images obtained by conventional transmission electron microscopy (TEM). The A-capsids are empty capsids that have lost the scaffold proteins (16, 17). The C-capsid is the mature capsid, which lacks the scaffold proteins and contains the viral genome (16, 17).

In the herpesviral lytic replication process, multiple copies of the herpesvirus genome are tandemly repeated, and one unit of the viral genome is cleaved at the TR as it is packaged into a capsid (18). KSHV has a TR of 801 base pairs (bp), which contains tandemly arranged GC-rich DNA (19). The single unit of the viral genome is packaged and cleaved by the viral terminase complex, which consists of three viral proteins. In KSHV, it has been hypothesized that ORF7, ORF29, and ORF67.5 comprise the terminase complex based on their sequence homology to other herpesviral terminase complex proteins. However, the details regarding the structural and functional properties of the KSHV terminase complex are largely unknown.

In HSV-1, UL15 (KSHV ORF29 homolog), UL28 (KSHV ORF7 homolog), and UL33 (KSHV ORF67.5 homolog) are components of the terminase complex (20, 21, 22). These HSV-1 proteins form a tripartite complex, and the hexameric ring of this tripartite complex becomes an HSV-1 terminase (22). UL15 and UL28 as well as UL28 and UL33 interact directly, but UL15 and UL33 do not interact in the absence of UL28 (23). If any of the components are defective, the viral genome is neither encapsidated nor cleaved, and capsid formation is arrested at the B-capsid stage (24, 25, 26, 27, 28, 29). Since these capsid formation pathways consist of virus-specific machineries that do not exist in any human cells, they may be targeted for the development of antiviral drugs. In fact, letermovir, a drug that targets the HCMV terminase complex, has been used clinically for HCMV treatment, and exhibits high antiviral activity and low adverse events (30, 31).

In contrast to HSV-1 and HCMV, little is known about the function of the KSHV terminase complex. ORF7, ORF29, and ORF67.5 are putative components of the KSHV terminase complex. We previously reported that there was no direct interaction between ORF29 and ORF67.5. However, we found that ORF7 interacted with both ORF29 and ORF67.5 (32). We also showed that ORF7-deficient KSHV formed numerous immature soccer ball-like capsids in the nucleus and failed to cleave the KSHV TR (33). The soccer ball-like capsid was a characteristic immature capsid that contains many particulate structures. Moreover, the soccer ball-like capsid was derived from procapsid (33). While these two reports indicate that ORF7 plays crucial roles in both KSHV terminase function and capsid formation, the function of ORF67.5 is completely unknown. Therefore, this study focused on the physiological function and virological significance of KSHV ORF67.5.

Other herpesviral homologs of KSHV ORF67.5 include HSV-1 UL33, HSV-2 UL33, VZV ORF25, EBV BFRF4, HCMV UL51, HHV-6 U35, and HHV-7 U35. KSHV ORF67.5 is 80 amino acids (aa) in length and is the shortest of the ORF67.5 homologs. HSV-1 UL33 is required for the cleavage of a single unit of the viral genome and its encapsidation. Interestingly, B-capsids accumulate in UL33-deficient HSV-1-infected cells (28, 29). VZV ORF25-deficient virus is unable to produce infectious virions (34). HCMV UL51 is also required for the cleavage of a single unit of the viral genome, and knockdown of HCMV UL51 results predominantly in B-capsid formation (35). There are no reports on the functions of HHV-6 U35 and HHV-7 U35. In addition, the function of EBV BFRF4, which belongs to the same gammaherpesvirinae subfamily as KSHV, is also unknown.

In this study, we generated ORF67.5-deficient KSHV and its revertant. These viruses were then used to analyze the function of ORF67.5. ORF67.5-deficient KSHV was unable to produce infectious virions, and soccer ball-like capsids were mainly observed in infected cells. Furthermore, the ORF67.5-deficient KSHV lacked the ability to cleave the TR. Moreover, ORF67.5 promoted the interaction between ORF7 and ORF29, and the interaction between ORF67.5 and ORF7 was also increased by ORF29.

## Results

### Construction of ORF67.5-deficient KSHV-BAC

In order to analyze the function of KSHV ORF67.5, we constructed ORF67.5-deficient KSHV-bacterial artificial chromosome (BAC) and revertant KSHV-BAC. The KSHV genome contains several ORFs with overlapping coding regions. The N-terminal coding region of ORF67.5 has been reported to overlap with the N-terminal coding region of the ORF68 gene, which encodes a total length of 545 aa (36). However, Glaunsinger et al. (37) did not detect endogenous ORF68 with 545 aa in KSHV-infected cells. Instead, they only detected the 467 aa form of ORF68, in which the N-terminal 78 aa region is deleted (37). Thus, the 545 aa ORF68 has been described as ORF68-Extended (37). The start codon (the first Met) of ORF68-Extended overlaps with the coding region of ORF67.5. In order to eliminate the effect of the ORF67.5 mutation on ORF68 expression, we deleted 1 bp upstream of the start codon in ORF68-Extended from wild-type KSHV-BAC to create ΔORF67.5-BAC16 (Fig. 1A). ΔORF67.5-BAC16 has a frameshift mutation (39^th^ G-C bp deletion) at the N-terminal side of the ORF67.5 coding region, by which a nonsense mRNA is transcribed. This nonsense mRNA contains 12 aa from the N-terminal side of the ORF67.5 coding region (80 aa) followed by 11 aa unrelated to ORF67.5 and a stop codon. In addition to ΔORF67.5-BAC16, Revertant-BAC16 was constructed by reinsertion of the 1 bp deletion into ΔORF67.5-BAC16 (Fig. 1A). The sequence of the mutagenesis site and neighboring nucleotides of the constructed BACs were confirmed by Sanger sequencing (Fig. 1A). The insertion and removal of the kanamycin-resistance cassette associated with the mutagenesis was confirmed by EcoRV digestion of each BAC followed by agarose gel electrophoresis (Fig. 1B).

**FIG 1.**
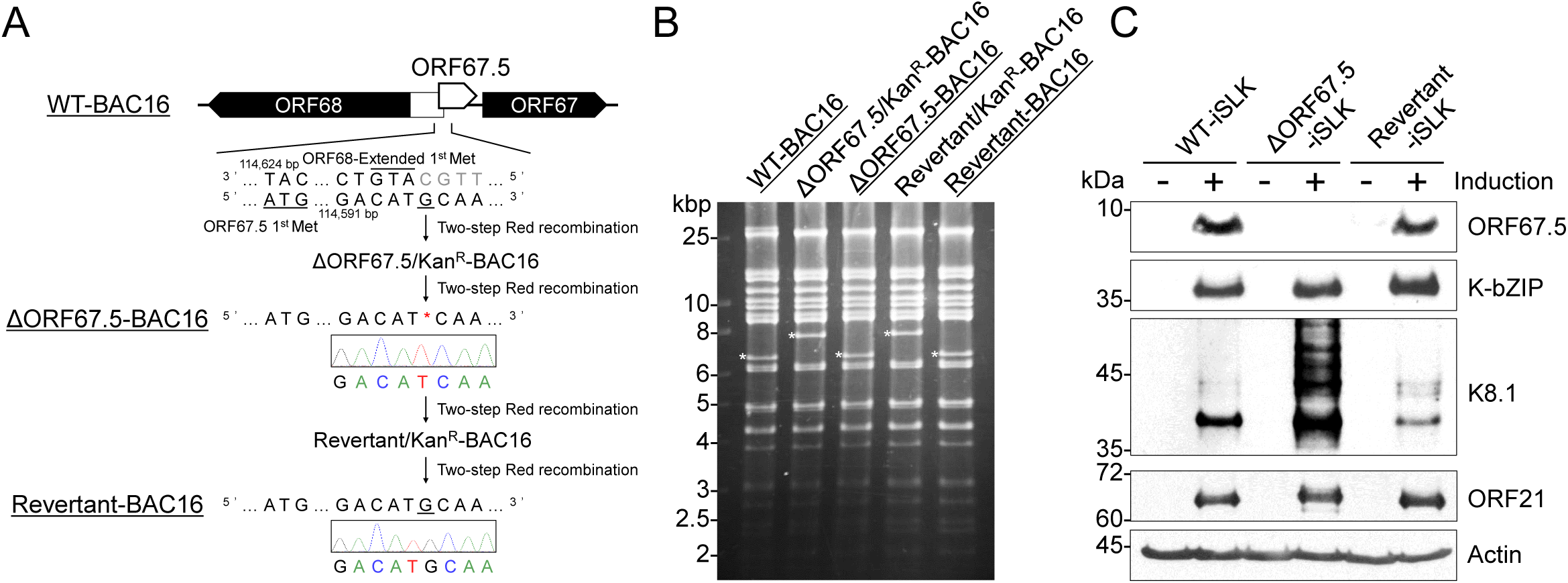
Construction of ΔORF67.5-BAC16 and Revertant-BAC16. (A) Diagram showing the location of KSHV ORF67.5 (nt 114382-114624; accession number: GQ994935) and its neighboring ORFs on the KSHV-BAC (WT-BAC16). The adjacent DNA sequence of the mutation sites in ΔORF67.5-BAC16 and Revertant-BAC16 clones was confirmed by Sanger sequencing. (B) Agarose gel electrophoresis image of EcoRV-digested recombinant BAC16 clones. The asterisks indicate the insertion or deletion of a kanamycin-resistance cassette in each BAC clone. (C) Western blot showing the disappearance of ORF67.5 expression in lytic-induced (+) or uninduced (−) ΔORF67.5-BAC16-harboring iSLK cells (ΔORF67.5-iSLK). WT-iSLK, ΔORF67.5-iSLK, and Revertant-iSLK were treated (or untreated) with Dox and SB for 72 h to induce lytic replication. An antibody against β-actin (Actin) was used as a loading control. The lytic gene products, K-bZIP (DE), K8.1 (L), and ORF21 (L) were expressed in all cell lines, unlike ORF67.5, which was absent in ΔORF67.5-iSLK.

Wild-type BAC16, ΔORF67.5-BAC16, and Revertant-BAC16 were transfected into iSLK cells, and transfectants were selected with hygromycin B to establish WT-iSLK, ΔORF67.5-iSLK, and Revertant-iSLK, cell lines, respectively. Hereafter, each cell line is simply referred to as WT-iSLK, ΔORF67.5-iSLK, and Revertant-iSLK. Treatment of iSLK cells with doxycycline (Dox) and sodium butyrate (SB) induces KSHV lytic infection (38). In order to confirm the disappearance of full-length ORF67.5 protein expression in ΔORF67.5-iSLK, we generated a rabbit polyclonal antibody (pAb) against ORF67.5. Each cell line was treated with Dox and SB to induce lytic infection, and the cells were lysed. Since ORF67.5 is a low molecular weight (MW) protein, Western blotting was performed using Tricine electrophoresis. In all cell lines, ORF67.5 expression was not detected when lytic infection was not induced. In contrast, when lytic infection was induced, ORF67.5 expression was confirmed in WT-iSLK and Revertant-iSLK, but not in ΔORF67.5-iSLK (Fig. 1C). The predicted MW of the endogenous ORF67.5 is 9.2 kDa. This Western blot using the generated anti-ORF67.5 antibody did not detect non-specific signals in the MW region below 15 kDa but did detect non-specific signals in the higher MW regions. We also examined the expression of lytic gene products other than ORF67.5 by Western blotting. The DE gene product K-bZIP and the L gene product ORF21 were detected to the same extent in all induced iSLK cell lines (Fig. 1C). In contrast, the L gene product K8.1 was upregulated in ΔORF67.5-iSLK compared to WT-iSLK and Revertant-iSLK (Fig. 1C). Although the reason for the increase in K8.1 expression is unknown, the same results were observed in multiple experiments. These data showed that ΔORF67.5-iSLK expressed the lytic gene products, K-bZIP, K8.1, and ORF21, but not ORF67.5.

### ORF67.5 is required for infectious virus production

To elucidate the virological functions of ORF67.5, we examined the contribution of ORF67.5 to lytic gene expression during lytic infection. Each iSLK cell line was treated with Dox and SB to induce lytic infection, and the transcription levels of ORF16 (IE), ORF46+ORF47 (DE), and K8.1 (L) were evaluated by reverse transcription-quantitative PCR (RT-qPCR). In all iSLK cell lines, we observed no remarkable differences in the mRNA expression levels of the tested lytic genes (Fig. 2A). In addition, qPCR measurement of the amount of intracellular viral genome replication showed no reduction of viral genomic DNA in ΔORF67.5-iSLK compared to WT-iSLK and Revertant-iSLK (Fig. 2B). We also analyzed the effect of ORF67.5 deficiency on virus production and infectious virus production. qPCR-mediated quantification of the amount of DNase-resistant KSHV genomic DNA in the culture supernatant of lytic-induced cells showed that ΔORF67.5-iSLK produced significantly less virus compared to WT-iSLK and Revertant-iSLK (Fig. 2C). Next, the infectivity of the progeny virus produced by ΔORF67.5-iSLK was analyzed. The culture supernatant of lytic-induced ΔORF67.5-iSLK was co-incubated with fresh HEK293T cells, and after 24 h, the percentage of green fluorescent protein (GFP)-positive HEK293T cells was measured by flow cytometry. Since BAC16 encodes the GFP gene, this supernatant transfer assay can assess the infectivity of the progeny virus by detecting GFP fluorescence in recipient cells (39). The infectivity of the progeny viruses produced in ΔORF67.5-iSLK was extremely low compared to those produced by WT-iSLK and Revertant-iSLK (Fig. 2D).

**FIG 2.**
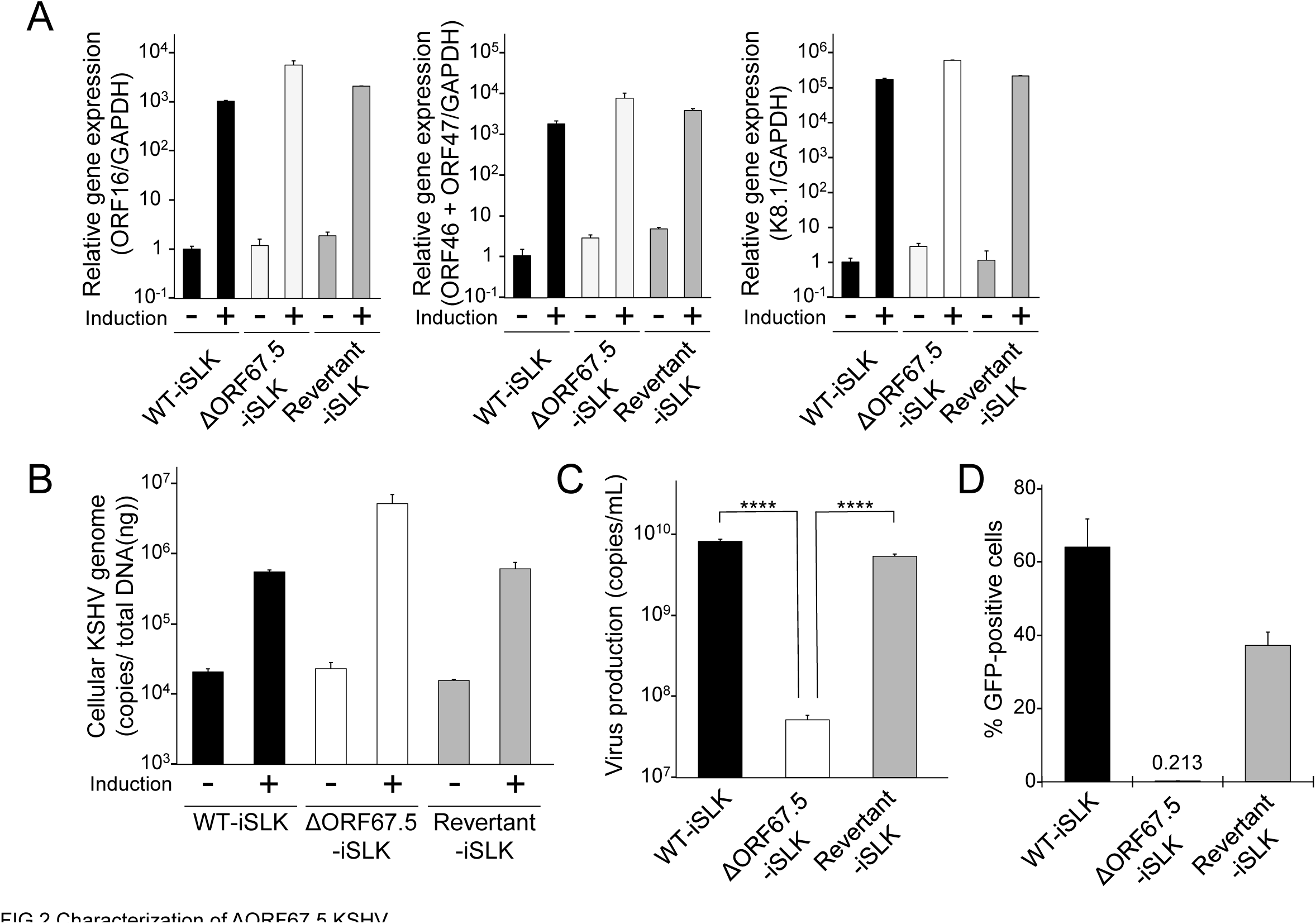
Characterization of ΔORF67.5 KSHV. (A) mRNA expression analysis of the following KSHV lytic genes: IE, ORF16; DE, ORF46 and ORF47; and L, K8.1. WT-iSLK, ΔORF67.5-iSLK, and Revertant-iSLK were treated with Dox and SB for 72 h to induce the lytic phase (+) or were uninduced (−). Total RNA was purified from harvested iSLK cells and subjected to RT-qPCR. The lytic gene mRNA levels were normalized to GAPDH mRNA levels. The values obtained from uninduced WT-iSLK were defined as 1.0. (B) Quantification of intracellular KSHV genomic DNA. WT-iSLK, ΔORF67.5-iSLK, and Revertant-iSLK were treated with Dox and SB to induce the lytic phase (+) or were uninduced (−). At 72 h post-induction, viral DNA was purified from the harvested cells. The intracellular viral genome copy number was then measured by qPCR and normalized to the amount of total DNA. (C) Quantification of extracellular encapsidated KSHV genomes. Each iSLK cell line was treated with Dox and SB for 72 h to induce the lytic phase, and the culture supernatants were harvested. Next, encapsidated KSHV genomic DNA from the culture supernatants was measured by qPCR. ****, P < 0.001. (D) Measurement of infectious virus. Each iSLK cell line was cultured with medium containing Dox and SB for 72 h to induce the lytic phase, and the culture supernatants were harvested. The supernatants were mixed with fresh HEK293T cells for infection with progeny virus. At 24 h post-infection, GFP-positive cells were counted by flow cytometry.

In order to further explore the involvement of ORF67.5 in virus production, we analyzed the complementation of virus production in ΔORF67.5-iSLK by exogenous ORF67.5 expression. ΔORF67.5-iSLK was transiently transfected with either empty plasmid or FLAG-tagged ORF67.5 expression plasmid and the transfectants were treated with Dox and SB to induce lytic infection. The amount of virus production in the culture supernatant was quantified by qPCR. WT-iSLK transfected with empty plasmid was used as a control. The results showed that the reduced virus production in ΔORF67.5-iSLK was significantly complemented by the expression of exogenous ORF67.5 (Fig. 3A). Expression of exogenous ORF67.5 was confirmed by Western blotting with an antibody against FLAG (Fig 3A, lower panel). In addition, we examined the complementation of infectious virus production in ΔORF67.5-iSLK by exogenous ORF67.5 expression. Empty plasmid-transfected WT-iSLK and either empty plasmid- or FLAG-ORF67.5-transfected ΔORF67.5-iSLK were treated with Dox and SB to induce lytic infection. Next, the infectivity of the progeny virus in the culture supernatant was evaluated by a supernatant transfer assay. We found that exogenous expression of ORF67.5 overcame the reduction in infectious virus production by the ORF67.5-deficient cells, ΔORF67.5-iSLK (Fig. 3B). These results indicate that ORF67.5 is dispensable for lytic gene transcription and viral genome replication, but it is required for virus production and infectious virus production.

**FIG 3.**
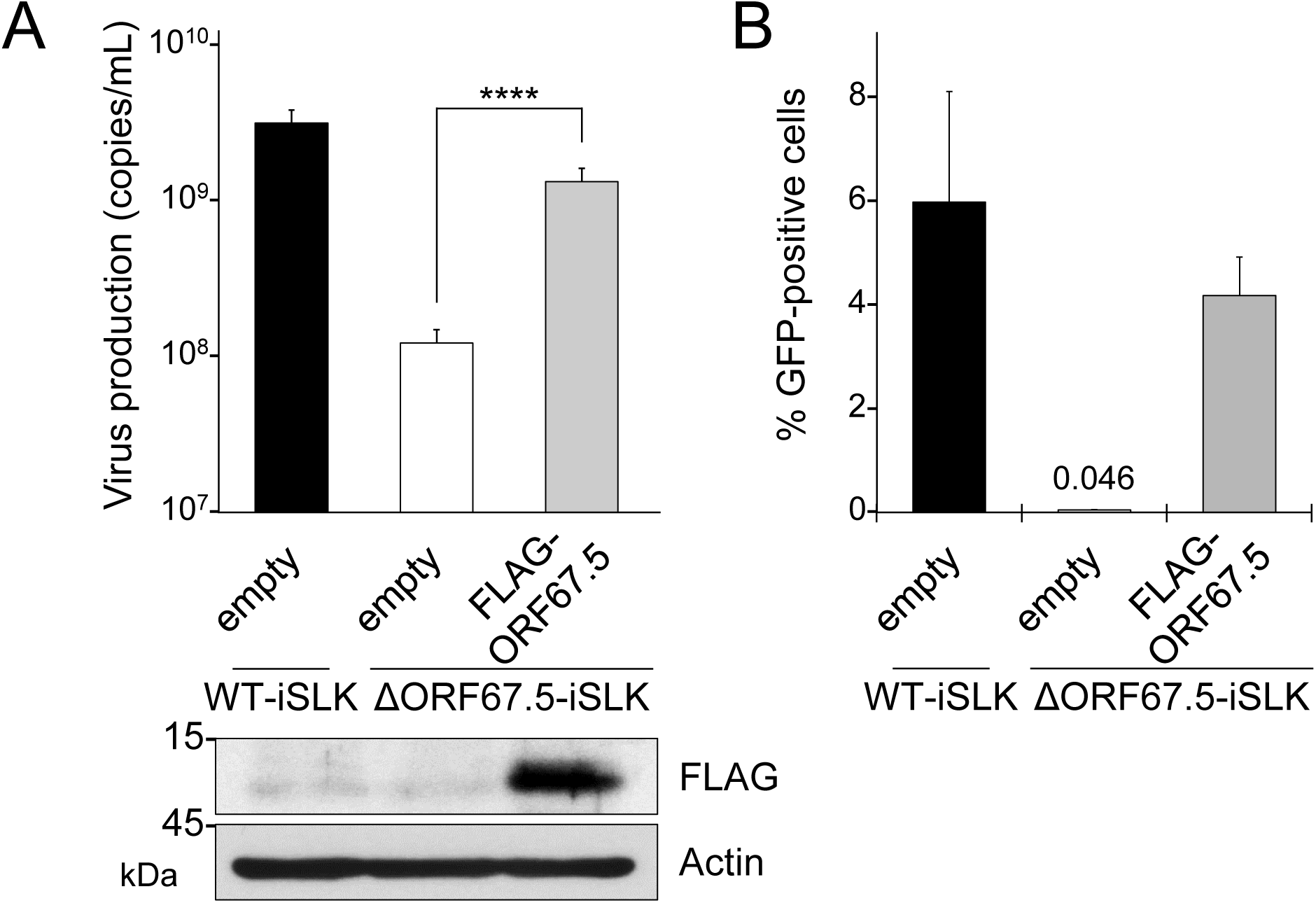
Complementation of ORF67.5-deficient KSHV with exogenous ORF67.5 expression. (A) Exogenous expression of ORF67.5 rescued the reduction in extracellular KSHV production observed in ΔORF67.5-iSLK. ΔORF67.5-iSLK was transiently transfected with empty plasmid (empty) or FLAG-ORF67.5 plasmid and was simultaneously cultured with medium containing Dox and SB for 72 h to induce the lytic phase. Encapsidated KSHV genomes in the culture supernatant were quantified by qPCR. Exogenous ORF67.5 expression was confirmed by Western blotting using an anti-FLAG (FLAG) antibody. An antibody against β-actin (Actin) was used as a loading control. ****, P < 0.001. (B) Exogenous expression of ORF67.5 rescued the reduction in infectious virus production observed in ΔORF67.5-iSLK. ΔORF67.5-iSLK was transiently transfected with empty plasmid (empty) or FLAG-ORF67.5 expression plasmid and simultaneously cultured with medium containing Dox and SB for 72 h to induce the lytic phase. Harvested culture supernatants were then used for infection of fresh HEK293T cells. At 24 h post-infection, GFP-positive cells were counted by flow cytometry.

### ORF67.5-deficient KSHV forms the soccer ball-like capsids

KSHV ORF7 and ORF67.5 can be presumed to be components of the KSHV terminase complex. We previously reported that in ORF7-deficient KSHV-harboring cells, most capsids formed in the nucleus during lytic infection were immature capsids, which resembled the telstar pattern of a soccer ball (33). Therefore, we referred to these capsids as “soccer ball-like capsids” (33). In this study, we analyzed capsid formation in lytic-induced ORF67.5-deficient KSHV-harboring cells. WT-iSLK and ΔORF67.5-iSLK were treated with Dox and SB for 48 h to induce lytic infection, and the morphology of capsids formed in the nuclei was observed by TEM. WT-iSLK produced all types of capsids, including A-capsids, B-capsids, C-capsids, and soccer ball-like capsids (Fig. 4A). In contrast, mature capsids (C-capsids) were not detected in ΔORF67.5-iSLK, and most of the capsids formed were soccer ball-like capsids (Fig. 4B). The quantification of each capsid type is shown in Fig. 4C. Our results indicate that ORF67.5 contributes to capsid formation, and deletion of ORF67.5 induces the formation of soccer ball-like capsids, which is also observed in lytically induced ORF7-deficient KSHV-harboring cells.

**FIG 4.**
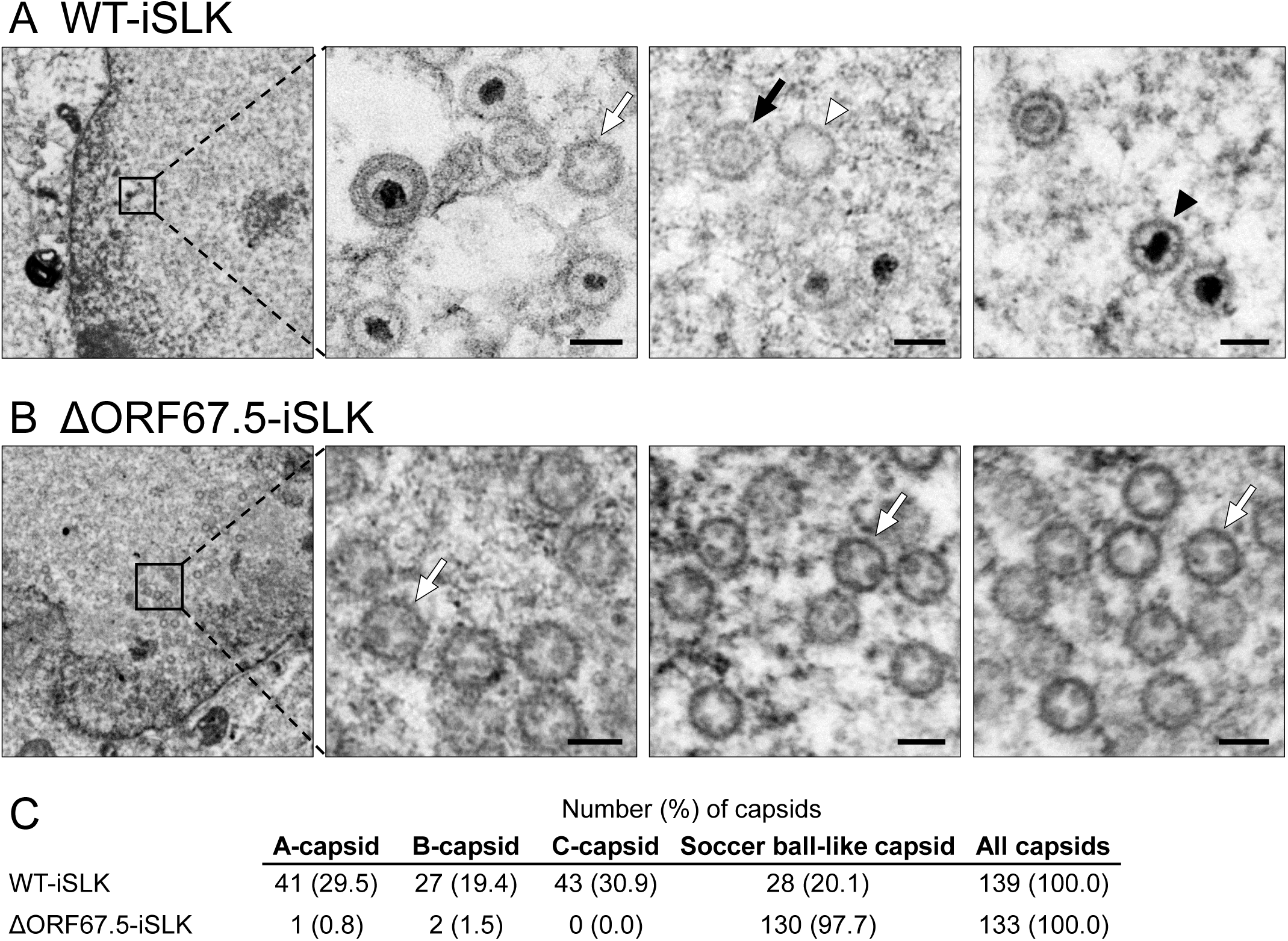
TEM images showing capsids formed in WT-iSLK and ΔORF67.5-iSLK. Cells were lytically induced for 48 h, and nuclear viral capsids in (A) WT-iSLK and (B) ΔORF67.5-iSLK were observed by TEM. The black arrowheads, white arrowheads, black arrows, and white arrows indicate C-capsids, A-capsids, B-capsids, and soccer ball-like capsids, respectively. The scale bars represent a length of 100 nm. (C) Quantification of each capsid type observed by TEM.

### ORF67.5 is required for TR cleavage of the KSHV genome

We analyzed the contribution of ORF67.5 in TR cleavage of the replicated KSHV genome during lytic infection. The cleaved TR fragments were detected by a method established by Glaunsinger et al. with some modifications (the same method by which we previously analyzed ΔORF7-KSHV) (33, 37). ΔORF67.5-iSLK was treated with Dox and SB for 72 h to induce lytic infection. Next, the total genomic DNA was digested with restriction enzymes and was subjected to Southern blotting using a probe against 1×TR to detect viral DNA fragments containing TRs. Uncleaved TR-containing DNA was detected in all cell lines and was increased in all lytic-induced cell lines compared to uninduced cell lines (Fig. 5). This data showed that KSHV genome replication in ΔORF67.5-iSLK was similar to that observed in WT-iSLK, supporting the results shown in Figure 2B (i.e., ORF67.5 is dispensable for viral genome replication) (Fig. 5). In contrast to the uncleaved TR, multiple cleaved TR fragments were detected in lytic-induced WT-iSLK and Revertant-iSLK, but not in lytic-induced ΔORF67.5-iSLK (Fig. 5). These fragment patterns were consistent with those in our previous report of ΔORF7-KSHV and in the report of ΔORF68-KSHV by Glaunsinger et al. (33, 37). These results demonstrate that ORF67.5 is required for TR cleavage of the KSHV genome.

**FIG 5.**
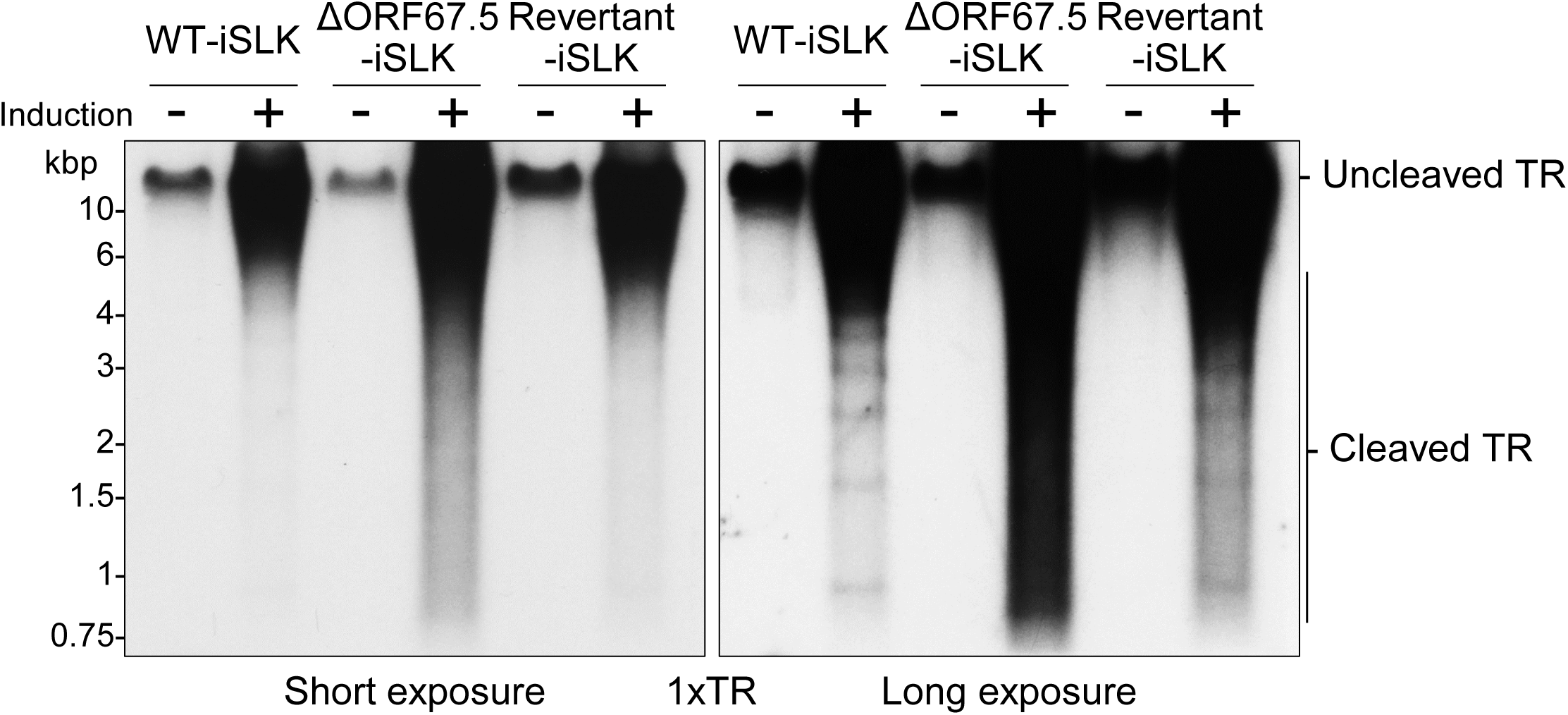
ΔORF67.5-KSHV failed to cleave the TR sites in the viral genome. WT-iSLK, ΔORF67.5-iSLK, and Revertant-iSLK were treated with (+) or without (−) Dox and SB for 72 h to induce the lytic phase. Purified total genomic DNA was digested with EcoRI and SalI and subjected to Southern blotting. Uncleaved and cleaved TRs were detected using a DIG-labeled 1×TR probe and their migration patterns are denoted to the right of the images. Left panel, short exposure; right panel, long exposure.

### The conserved regions of ORF67.5 are important for virus production

To obtain further information on the functional significance of the KSHV ORF67.5 protein, we aligned the aa sequences of the human herpesviral ORF67.5 homologs. Figure 6A shows the aa sequence alignments of KSHV ORF67.5 homologs from HSV-1, HSV-2, VZV, EBV, HCMV, HHV-6, and HHV-7. The aa residues that completely match are shown in white letters with a black background and similar aa residues are shown in black letters with a gray background. This analysis revealed the conserved regions within the KSHV ORF67.5 protein (Fig. 6A). Interestingly, KSHV ORF67.5 has the shortest aa sequence among its homologs. In order to determine the functionally important region(s) of ORF67.5 that contribute to virus production, we generated a total of 13 alanine mutants spanning the conserved regions (m2-6 and m8-12) and non-conserved regions (m1, m7, and m13) of ORF67.5 (Fig. 6B). In this report, each mutated region in ORF67.5 and each ORF67.5 alanine mutant are referred to as m1-13 and ORF67.5 m1-13, respectively (Fig. 6).

**FIG 6.**
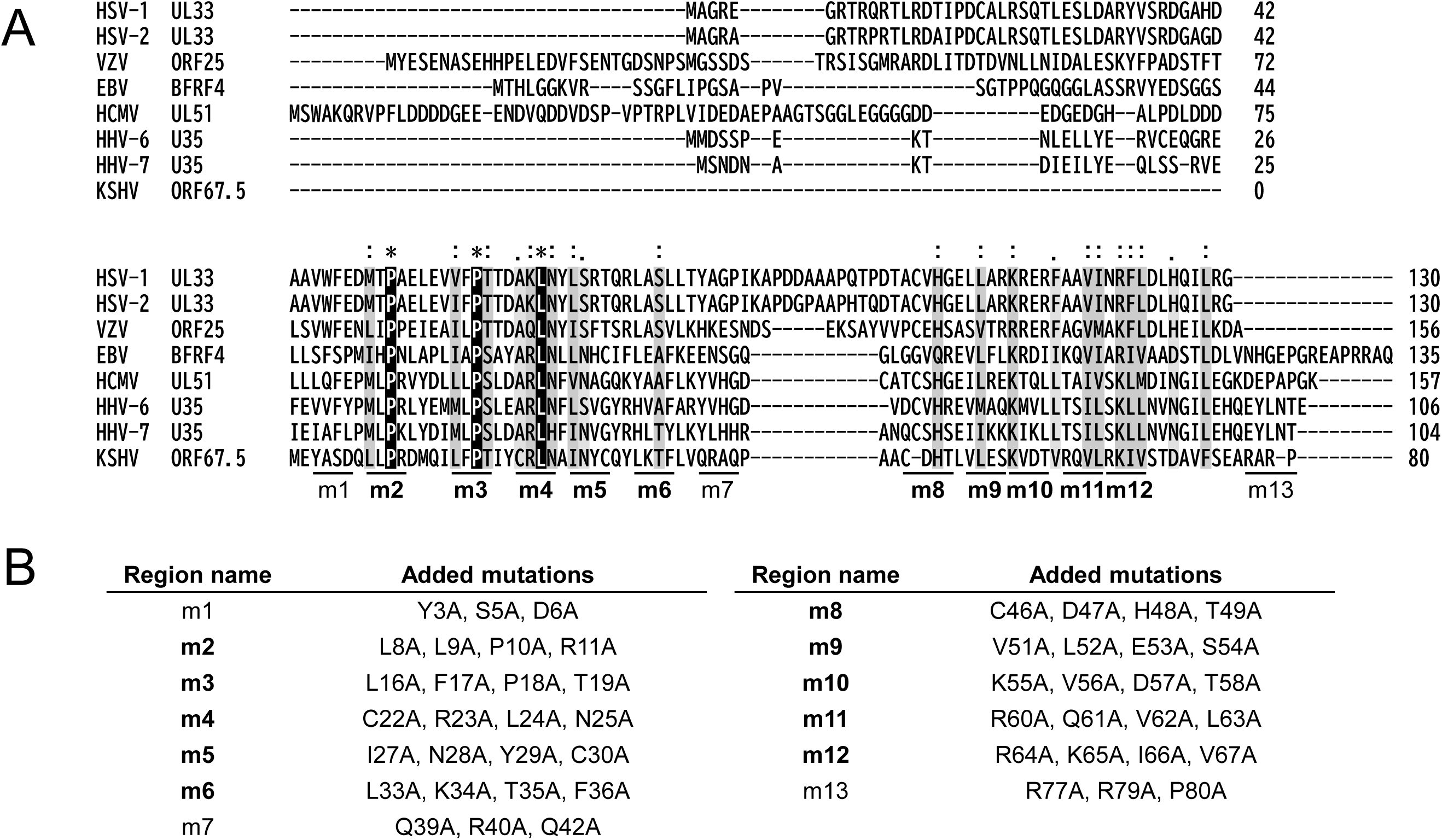
Amino acid sequence alignment between KSHV ORF67.5 and its human herpesviral homologs as well as construction of the ORF67.5 alanine mutants. (A) Amino acid sequences were obtained from the NCBI database [HSV-1 UL33 (GenBank: QAU10292), HSV-2 UL33 (NCBI Reference Sequence: YP_009137185), VZV ORF25 (GenBank: QXM18354), EBV BFRF4 (GenBank: AAA45868), HCMV UL51 (GenBank: QHB20503), HHV-6 U35 (NCBI Reference Sequence: NP_042928), HHV-7 U35 (GenBank: AAC40749), and KSHV ORF67. 5 (GenBank: QFU18781)]. The raw data used for the alignment were obtained with the Clustal Omega program (EMBL-EBI; https://www.ebi.ac.uk/Tools/msa/clustalo/). The background indicates the degree of homology. The completely matched aa residues are shown in white letters with a black background and similar aa residues are shown in black letters with a gray background. The conserved regions (m2-6 and m8-12) and the non-conserved regions (m1, m7, and m13) among the human herpesviral ORF67.5 homologs are indicated at the bottom of the sequence alignments and the conserved regions are indicated in bold. (B) The aa sequences of the conserved and non-conserved regions of the human herpesviral ORF67.5 homologs that were substituted with alanine in each ORF67.5 mutant. The names of the conserved regions are indicated in bold.

We then used these mutants to analyze the impact of the KSHV ORF67.5 conserved regions on virus production (Fig. 7). To this end, we conducted a complementation assay to determine if any of the ORF67.5 alanine mutant (m1-13) expression plasmids could rescue virus production in ΔORF67.5-iSLK. ΔORF67.5-iSLK was transfected with each alanine mutant plasmid, and lytic infection was induced with Dox and SB. The culture supernatants were harvested 72 h after induction, and virus production in the supernatants was measured by qPCR. The low virus production in ΔORF67.5-iSLK was complemented by transfection of the ORF67.5 wild-type (WT) expression plasmid (Fig. 7, upper panel), which was also shown in Figure 3A. On the other hand, among the ORF67.5 alanine mutants, only ORF67.5 m1, m7, and m13 restored the reduction in virus production from ΔORF67.5-iSLK, while the other mutants failed to restore it (Fig. 7, upper panel). The expression of each ORF67.5 mutant was confirmed by Western blotting (Fig. 7, lower panel). These results indicate that the m2-6 and m8-12 regions in ORF67.5 are essential for the function of ORF67.5 in virus production, whereas the m1, m7, and m13 regions in ORF67.5 are not essential for the function of ORF67.5 in virus production. Importantly, the m2-6 and m8-12 regions within the ORF67.5 protein are highly conserved among herpesviruses, whereas the m1, m7, and m13 regions are not conserved.

**FIG 7.**
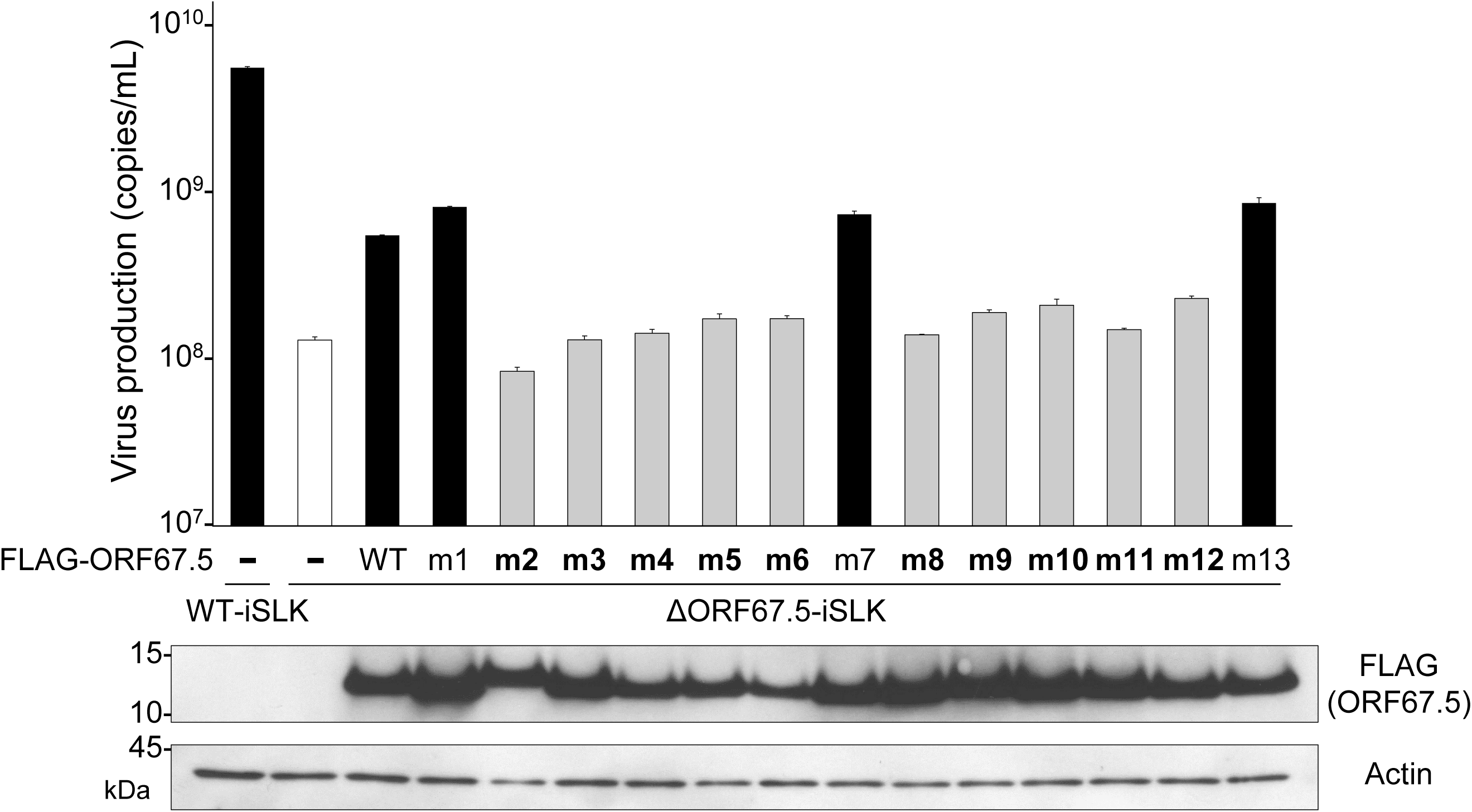
The ORF67.5 conserved region alanine mutants failed to rescue the reduction in virus production from ΔORF67.5-iSLK. In order to evaluate the contribution of the conserved and non-conserved regions of ORF67.5 to virus production, complementation assays were performed by exogenous ORF67.5 alanine mutant expression in ΔORF67.5-iSLK. ΔORF67.5-iSLK was transiently transfected with FLAG-ORF67.5 WT or FLAG-ORF67.5 alanine mutant expression plasmids (FLAG-ORF67.5 m1-m13). During transfection, the cells were simultaneously cultured in medium containing Dox and SB for 72 h to induce the lytic phase. Encapsidated KSHV genomes in the culture supernatants were quantified by qPCR. The expression of exogenous FLAG-tagged ORF67.5 WT and mutants were confirmed by Western blotting using an anti-FLAG (FLAG) antibody. An antibody against β-actin (Actin) was used as a loading control.

### The conserved regions of ORF67.5 are important for its interaction with ORF7

We previously reported that a pull-down assay did not detect a direct interaction between ORF67.5 and ORF29, but a direct interaction between ORF67.5 and ORF7 was detected (32). Furthermore, we showed above that the m2-6 and m8-12 regions in the ORF67.5 protein, which are conserved among herpesviral ORF67.5 homologs, are important for the virus-producing function of ORF67.5 (Figs. 6 and 7). Therefore, we tested whether the conserved regions in ORF67.5 contribute to its direct interaction with ORF7. To this end, pull-down assays were performed using the ORF67.5 m1-13 alanine mutants. Each FLAG-tagged ORF67.5 alanine mutant plasmid was co-transfected together with an S-tagged ORF7 plasmid into HEK293T cells. Next, S-tagged ORF7 was precipitated with S-protein agarose to detect the interaction between ORF7 and each ORF67.5 mutant. Our results showed that ORF67.5 WT interacted with ORF7, and the ORF67.5 m1, m7, and m13 mutants also interacted with ORF7 (Fig. 8). However, the ORF67.5 m2-6 and m8-12 mutants failed to interact with ORF7 (Fig. 8). These data indicate that the conserved regions (m2-6 and m8-12) in the ORF67.5 protein are involved in both virus production and the interaction of ORF67.5 with ORF7.

**FIG 8.**
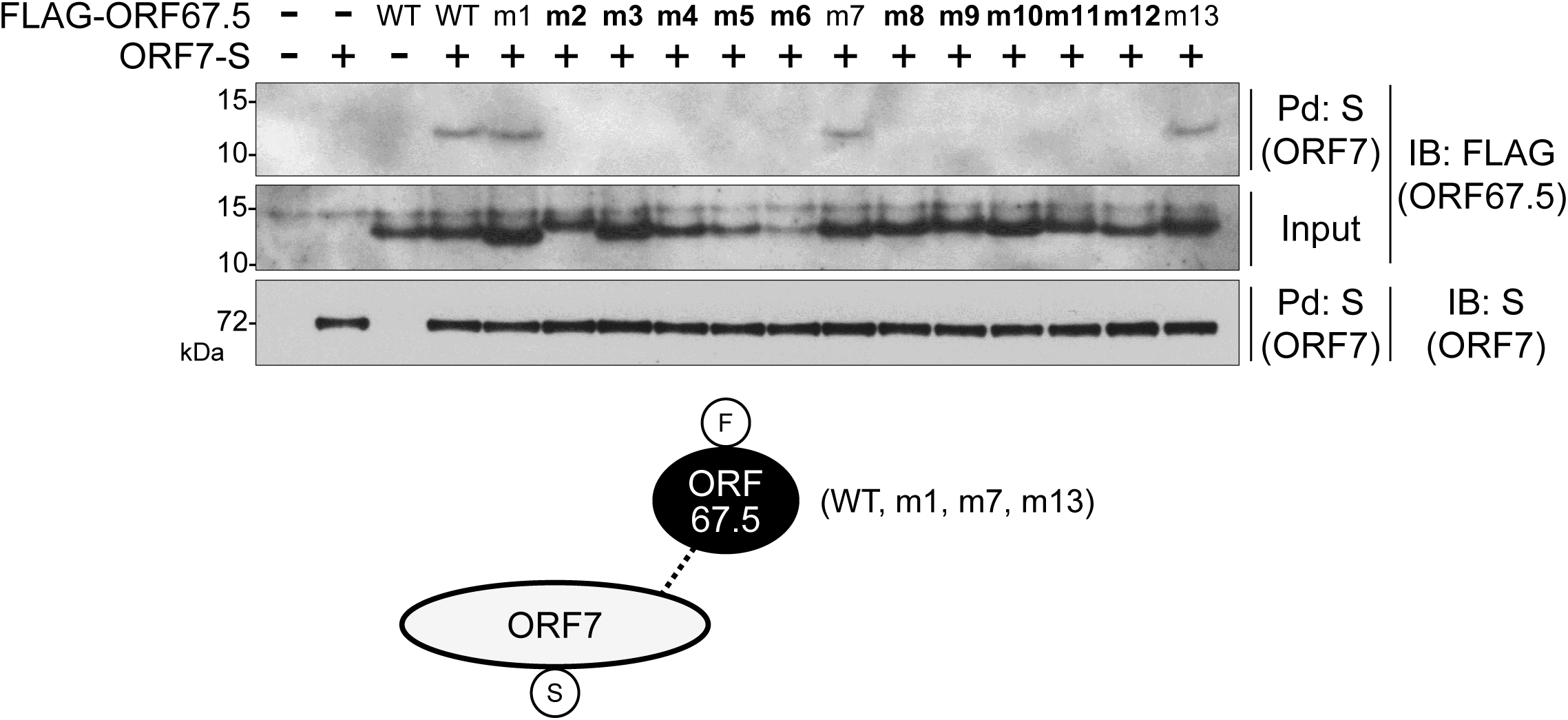
The conserved regions of ORF67.5 were necessary for the interaction of ORF67.5 with ORF7. An expression plasmid encoding FLAG-tagged wild-type ORF67.5 (FLAG-ORF67.5 WT) or FLAG-tagged ORF67.5 mutant (FLAG-ORF67.5 m1-m13) was co-transfected with a S-tagged ORF7 expression plasmid (ORF7-S) into HEK293T cells. Next, the cell lysates were subjected to pull-down assays (Pd) using S-protein immobilized beads (S) for the precipitation of ORF7-S protein. Any FLAG-ORF67.5 protein that interacted with ORF7 was detected by Western blotting using an anti-FLAG antibody (FLAG). The proteins in parenthesis indicate which protein was recognized by the assay. The conserved regions of ORF67.5 are indicated in bold. The bottom panel shows a schematic representation of the results with the protein tags indicated in the circles.

### ORF67.5 promotes the interaction between ORF7 and ORF29, and the interaction between ORF67.5 and ORF7 is increased by ORF29

ORF7 has been reported to interact with both ORF29 and ORF67.5, although a direct interaction between ORF67.5 and ORF29 was not detected in pull-down assays (32). Therefore, we investigated the effect of ORF67.5 on the interaction between ORF7 and ORF29. When FLAG-tagged ORF29 and S-tagged ORF7 were co-transfected into HEK293T cells, ORF7 interacted with ORF29 (Fig. 9A). Remarkably, co-transfection of HA-tagged ORF67.5 increased the interaction between ORF7 and ORF29 in an ORF67.5 expression level-dependent manner (Fig. 9A). The Western blotting data of the input samples showed that the presence of ORF67.5 had no effect on the ORF7 nor ORF29 expression levels (Fig. 9A). These results revealed that ORF67.5 promotes the interaction between ORF7 and ORF29.

**FIG 9.**
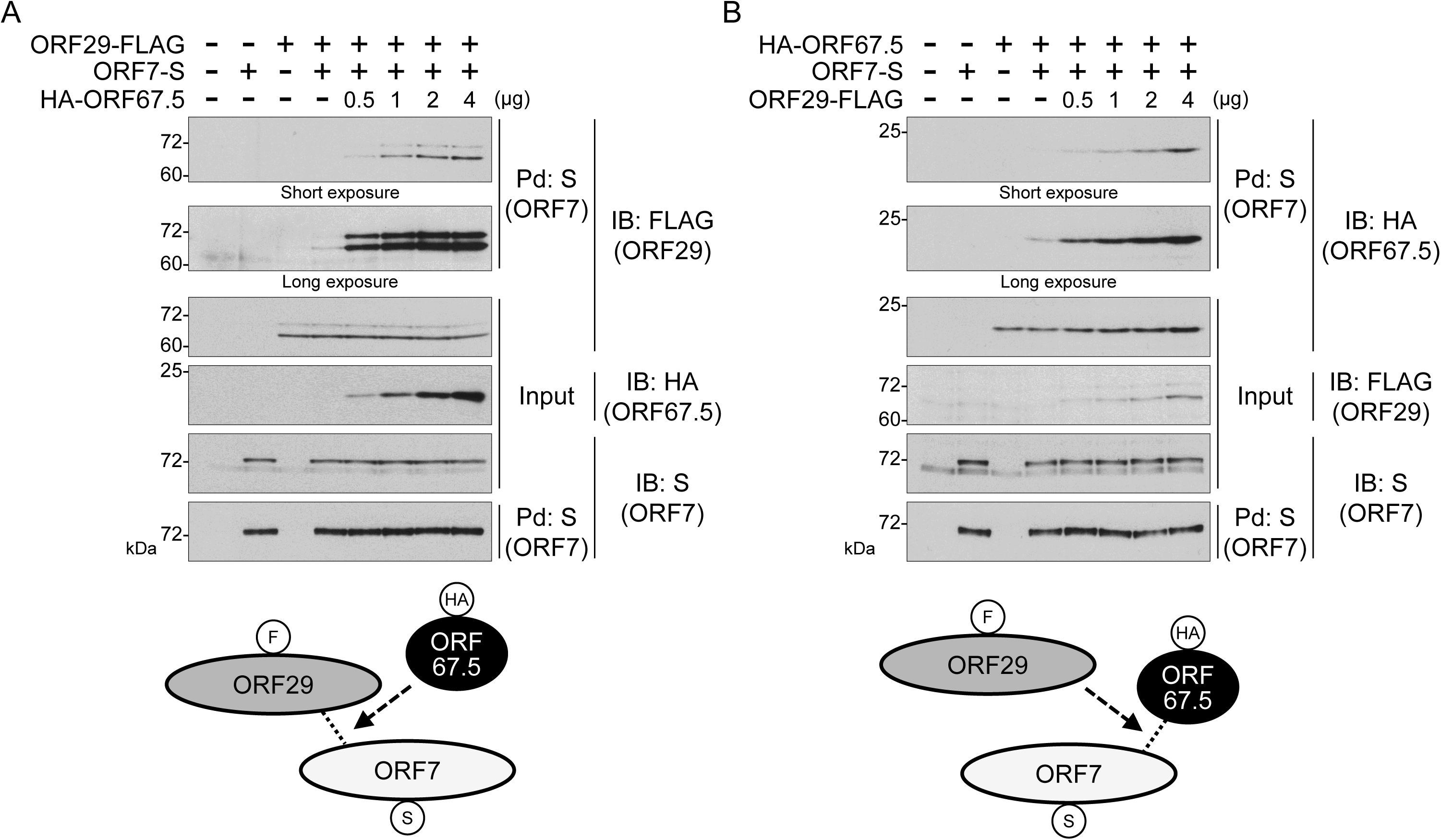
ORF67.5 promoted the interaction between ORF7 and ORF29, and ORF29 increased the interaction between ORF67.5 and ORF7. (A) Expression plasmids encoding ORF29-FLAG and ORF7-S were co-transfected with varying amounts of HA-ORF67.5 expression plasmid into HEK293T cells. Next, the cell lysates were subjected to pull-down (Pd) assays using S-protein immobilized beads (S) for the precipitation of ORF7-S protein. Any ORF29-FLAG protein that interacted with ORF7 was detected by Western blotting with an anti-FLAG antibody (FLAG). The proteins in parenthesis indicate which protein was recognized by the assay. The bottom panel shows a schematic representation of the results with the protein tags indicated in the circles. (B) Expression plasmids encoding HA-ORF67.5 and ORF7-S were co-transfected with varying amounts of ORF29-FLAG expression plasmid into HEK293T cells. Next, the cell lysates were subjected to pull-down (Pd) assays using S-protein immobilized beads (S), followed by Western blotting with anti-HA antibody (HA) to detect the ORF7-binding HA-ORF67.5 protein. The proteins in parenthesis indicate which protein was recognized by the assay. The bottom panel shows a schematic representation of the results with the protein tags indicated in the circles.

Next, we examined the effect of ORF29 on the interaction between ORF7 and ORF67.5. When HA-tagged ORF67.5 and S-tagged ORF7 were co-transfected into HEK293T cells, ORF7 interacted with ORF67.5 (Fig. 9B). As expected, co-transfection of FLAG-tagged ORF29 increased the interaction between ORF7 and ORF67.5 in an ORF29 expression level-dependent manner (Fig. 9B). The Western blotting data of the input samples showed that the presence of ORF29 had no effect on the ORF7 nor ORF67.5 expression levels (Fig. 9B). These results showed that the interaction between ORF67.5 and ORF7 is increased by ORF29.

### Identification of the regions in ORF67.5 responsible for promoting the interaction between ORF7 and ORF29

We disclosed that the conserved region mutants (m2-6 and m8-12) of ORF67.5 failed to rescue the reduction in virus production from ΔORF67.5-iSLK (Fig. 7) and failed to interact with ORF7 (Fig. 8). In other words, these regions, which are highly conserved among human herpesvirus homologs, are important for virus production and the interaction between ORF67.5 and ORF7. In addition, we found that the interaction between ORF7 and ORF29 was increased by ORF67.5, and the interaction between ORF7 and ORF67.5 was increased by ORF29 (Fig. 9). Therefore, we investigated whether the interaction between ORF7 and ORF29 was increased by each ORF67.5 mutant. Moreover, we determined whether the interaction between ORF7 and each ORF67.5 mutant was increased by ORF29 (Fig. 10A). As expected, the interaction between ORF7 and ORF29 was increased by the wild-type ORF67.5 (WT) and the non-conserved region mutants (ORF67.5 m1, m7, and m13), which rescued virus production in ΔORF67.5-iSLK as shown in Figure 7. The interaction of ORF7 with the non-conserved region mutants (ORF67.5 m1, m7, and m13) was also increased by ORF29 (Fig. 10A). Interestingly, in addition to the non-conserved region mutants (ORF67.5 m1, m7, and m13), the conserved region mutants (ORF67.5 m9, m11, and m12) also increased the interaction between ORF7 and ORF29. Furthermore, the interaction of ORF7 with the conserved region mutants (ORF67.5 m9, m11, and m12) was increased by ORF29 (Fig. 10A). However, these conserved region mutants (ORF67.5 m9, m11, and m12) could not overcome the reduction in virus production from ΔORF67.5-iSLK (Fig. 7) and did not have the ability to interact with ORF7 in the absence of ORF29 (Fig. 8). Moreover, the conserved region mutants (ORF67.5 m2, m3, m4, m5, m6, m8, and m10) failed to interact with ORF7 neither in the absence (Fig. 8) nor in the presence (Fig. 10A) of ORF29 and failed to increase the interaction between ORF7 and ORF29. Based on these findings, the non-conserved regions (m1, m7, and m13) of ORF67.5 had no effect on virus production nor the promotion of the interaction between ORF7 and ORF29. In contrast, the conserved regions (m9, m11, and m12) of ORF67.5 were required for its effects on virus production and for the interaction of ORF67.5 with ORF7 in the absence of ORF29. However, these conserved m9, m11, and m12 regions of ORF67.5 were not required for the interaction of ORF67.5 with ORF7 in the presence of ORF29, nor for the ORF67.5-mediated promotion of the interaction between ORF7 and ORF29. Since the conserved regions [m2 (L8, L9, P10, R11), m3 (L16, F17, P18, T19), m4 (C22, R23, L24, N25), m5 (I27, N28, Y29, C30), m6 (L33, K34, T35, F36), m8 (C46, D47, H48, T49), and m10 (K55, V56, D57, T58)] of ORF67.5 contribute to both virus production and the interaction of ORF67.5 with ORF7 (regardless of the presence or absence of ORF29), these regions are critically important for ORF67.5 function.

**FIG 10.**
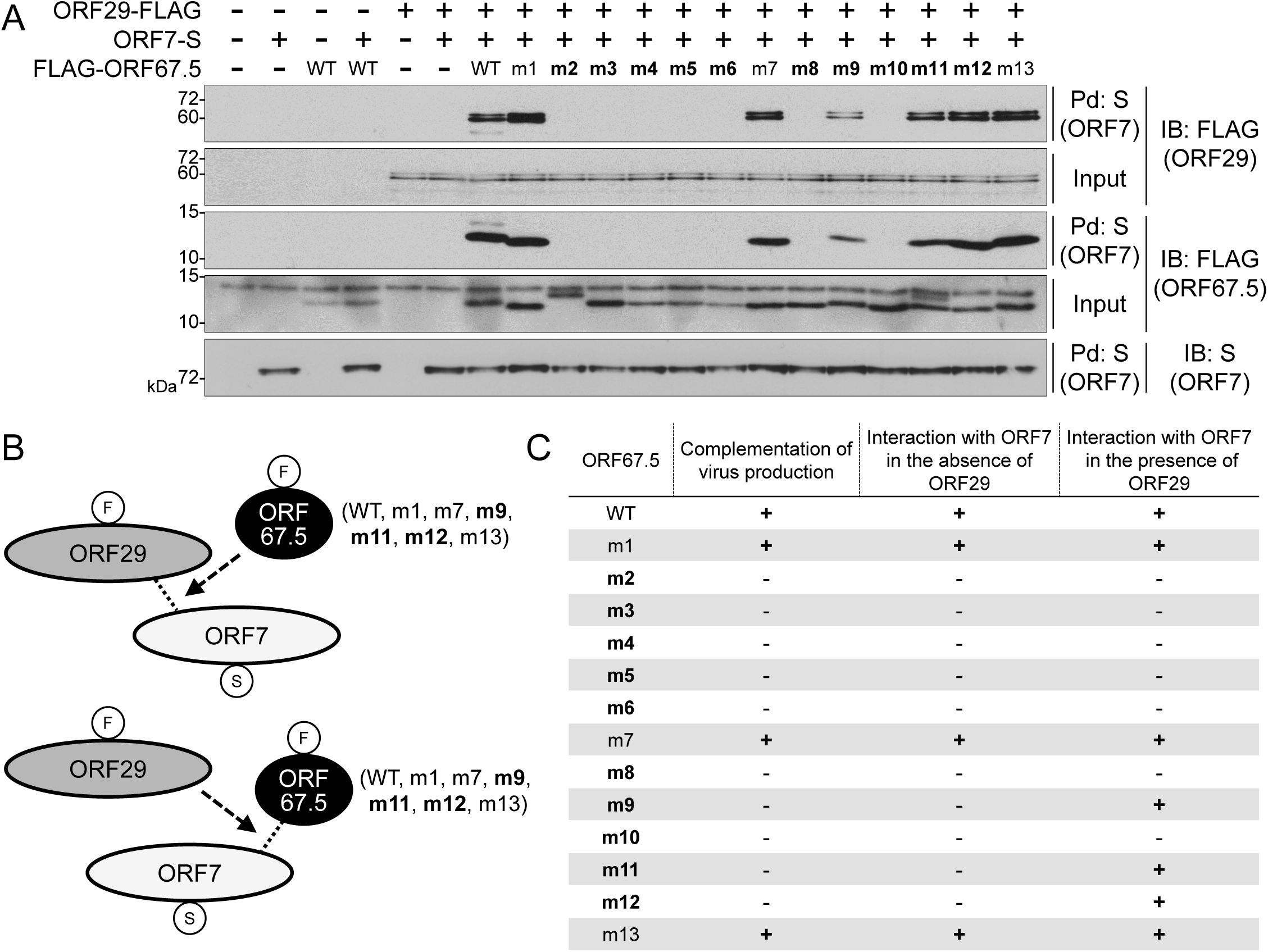
The effects of the conserved and non-conserved region mutants of ORF67.5 on the interaction between ORF7 and ORF29, as well as the effect of ORF29 on the interaction between each ORF67.5 mutant and ORF7. (A) Expression plasmids encoding ORF29-FLAG and ORF7-S were co-transfected with the FLAG-ORF67.5 mutant expression plasmid into HEK293T cells. Next, the cell lysates were subjected to pull-down (Pd) assays using S-protein immobilized beads (S) followed by Western blotting using anti-FLAG or anti-S antibodies (FLAG and S, respectively). Any ORF29-FLAG protein that interacted with ORF7-S in the presence of wild-type (WT) or alanine mutant ORF67.5 (m1-m13) was detected by Western blotting with anti-FLAG antibody. The proteins in parenthesis indicate which protein was recognized by the assay. (B) Schematic representation of the interaction relationship among ORF7, ORF29, and ORF67.5 in this experiment. The protein tags are indicated in the circles and the conserved regions of ORF67.5 are shown in bold. (C) A table summarizing the characteristics of the wild-type (WT) and alanine mutants of ORF67.5 (m1-m13) revealed by our experiments. The table summarizes the results obtained in Figure 7 (complementation of virus production), Figure 8 (the interaction with ORF7 in the absence of ORF29), and Figure 10A (the interaction with ORF7 in the presence of ORF29). The conserved regions in ORF67.5 are indicated in bold.

## Discussion

In this study, we showed that KSHV ORF67.5 was essential for infectious virus production, normal capsid formation, and TR cleavage of viral genomes. In addition, ORF67.5 enhanced the interaction between ORF7 and ORF29, and ORF29 also increased the interaction between ORF67.5 and ORF7. These results suggest that the KSHV terminase complex is composed of ORF7, ORF29, and ORF67.5, which are the KSHV homologs of the HSV-1 terminase complex components, UL28, UL15, and UL33, respectively. ORF67.5 facilitates the formation of the ORF7-ORF29-ORF67.5 complex and is crucial for the functional activity of the terminase complex, including TR cleavage. In addition, the conserved regions (m2-6 and m8-12) of ORF67.5 are important for virus production. Additionally, m2-6, m8, and m10 are required not only for virus production but also for the ORF67.5-mediated promotion of the interaction between ORF7 and ORF29. To our knowledge, this is the first report describing the virological significance and functions of ORF67.5 as a component of the KSHV terminase complex.

Numerous soccer ball-like capsids were observed in lytic-induced ΔORF67.5-iSLK (Fig. 4B). Capsids similar in morphology to the soccer ball-like capsids have also been observed in VZV and KSHV (33, 40, 41, 42, 43, 44). The soccer ball-like capsid is thought to be a state in which degraded scaffold proteins remain in the capsid after procapsid formation (33, 40). Similar to KSHV ORF67.5, deletion of KSHV ORF7, which is hypothesized to be one of the components of the KSHV terminase complex, also resulted in the formation of soccer ball-like capsids (33). The HSV-1 terminase complex excises and packages one unit of the viral genome from serially replicated viral genome precursors. During lytic replication, soccer ball-like capsid formation occurs when the ORF7- or ORF67.5-defective KSHV-harboring cell exhibits a defect in TR cleavage of the viral genome. These findings suggest that the KSHV capsid formation process may be arrested at the soccer ball-like capsid stage when viral genome precursor cleavage or genome packaging fails. During the KSHV capsid maturation process, when the inner scaffold layer of the procapsid is detached from the outer capsid by the viral protease ORF17, the degradation and disruption of the scaffold inner structure is proceeded (45). If the soccer ball-like capsids contain residual products derived from the disrupted inner scaffold structure, the terminase complex would not be required for the disruption (or degradation) of the scaffold proteins within the capsid. However, the external release of the disrupted scaffold structure may either require an unknown novel function of the KSHV terminase complex or insertion of the KSHV genome into the capsid. In other words, the collapsed scaffold proteins may be eliminated from the capsid through gaps in the outer capsid layer when these residual scaffold proteins are pushed out by the insertion of the KSHV genome. This hypothesis has also been proposed by Zhou et al. (40). Based on this hypothesis, when ORF7 or ORF67.5 is defective, the insertion of the KSHV genome into the capsid does not occur and the collapsed scaffold inner structures are not eliminated from the capsid. Thus, the capsid formation process may be arrested at the soccer ball-like capsid stage. When herpesvirus genome insertion occurs, the state of the capsid is still unclear, but further development of observational techniques such as time-lapse cryo-electron microscopy (cryo-EM) will be required to confirm the state in real time (11, 46).

KSHV ORF67.5 has the shortest aa sequence among its human herpesviral homologs (Fig. 6A). The KSHV ORF67.5 homolog, HSV-1 UL33, has an additional 42 aa on the N-terminal side, which is not present in KSHV ORF67.5. The N-terminal region of HSV-1 UL33 does not contribute to virus production, and its biological function is unknown (47). The m2-6 and m8-12 regions are highly conserved among the human herpesviral KSHV ORF67.5 homologs (Fig. 6A). The conserved region mutants of KSHV ORF67.5 (ORF67.5 m2-6 and m8-12) could not complement the reduction in virus production from ΔORF67.5-iSLK (Fig. 7). Further, we did not detect the interaction of ORF7 with these mutants in the absence of ORF29 (Fig. 8). In contrast, the non-conserved region mutants of KSHV ORF67.5 (ORF67.5 m1, m7, and m13), complemented virus production (Fig. 7) and interacted with ORF7 (Fig. 8). These findings suggest that the important regions within ORF67.5 that are involved in the formation and function of the KSHV terminase complex are highly conserved among human herpesviral homologs.

To further discuss the functionally critical regions of KSHV ORF67.5, the conformation of the tripartite complex comprising ORF7, ORF29, and ORF67.5 was predicted using the AI deep learning algorithm AlphaFold2 (Fig. 11). The conformation of the HSV-1 terminase complex composed of HSV-1 UL28 (green), UL15 (light blue), and UL33 (light purple) has already been determined using cryo-EM (22). We compared this conformation with the AI-predicted tripartite complex conformation of ORF7 (green), ORF29 (light blue), and ORF67.5 (light purple) (Figs. 11A and B). The overall conformation was largely consistent between the HSV-1 terminase complex and the putative KSHV terminase complex, with several differences. In the KSHV ORF7, ORF29, and ORF67.5 tripartite complex, ORF67.5 appeared to be wrapped in ORF7 and was not associated with ORF29. From another point of view, the ORF7-interacting ORF67.5 resembled a wheel clamp that supported the conformation of ORF7. This AI-predicted conformational data generally supported the results obtained by the pull-down assay (i.e., ORF67.5 interacted with ORF7, but not with ORF29) (32). It should be noted that in the actual HSV-1 terminase complex, tripartite complexes composed of UL28, UL15, and UL33 form a hexameric ring structure (22). KSHV ORF67.5 has three intramolecular helical structures, and its N- and C-terminal regions were hypothesized to fluctuate freely and not form a characteristic secondary structure (Fig. 11B). The m1-13 regions of ORF67.5 analyzed in this paper are indicated in this prediction model. In ORF67.5, the non-conserved region m7 is located at the junction of the helical structure, and the non-conserved regions m1 and m13 are located at the N-terminal and C-terminal regions which were hypothesized to fluctuate freely (Fig. 11C). The conserved regions m2-6 and m8-12 are all located in the helical structure of ORF67.5 (Fig. 11C). These models suggest that the formation of the helical structures is likely important for ORF67.5 function and may explain why the aa sequences of the helical structure-forming regions are highly conserved. This explanation is supported by our data, showing that mutations in the conserved regions (m2-6 and m8-12) of ORF67.5 impaired virus production (Fig. 7). This structural model is a prediction model, and the three-dimensional structure of the actual KSHV terminase complex needs to be elucidated by cryo-EM or X-ray structure analysis.

**FIG 11.**
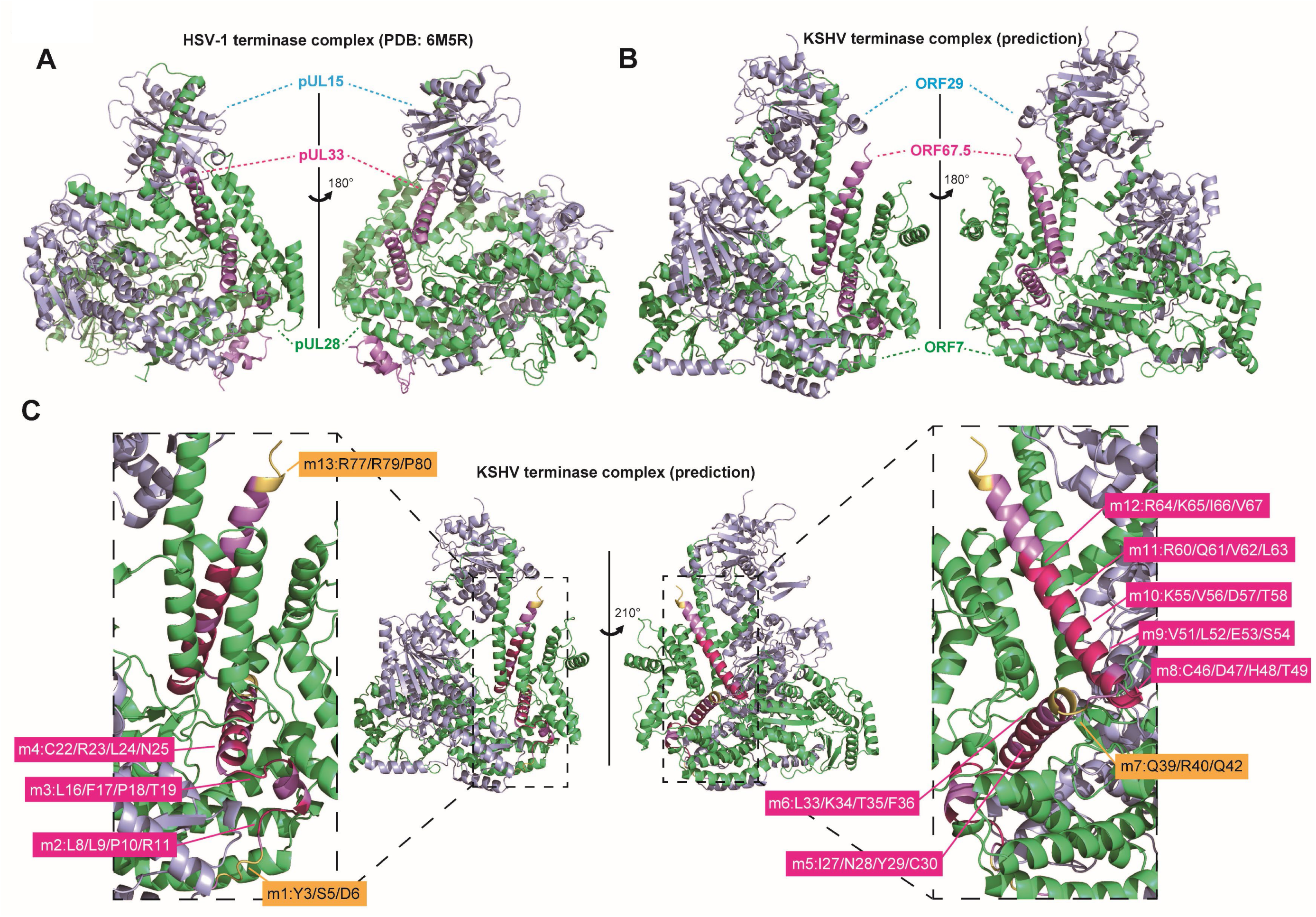
The structure of the HSV-1 terminase complex and the predicted structure model of the KSHV terminase complex. (A) The HSV-1 monomeric terminase complex structure (PDB; 6M5R) (22). Each HSV-1 protein cartoon was colored as follows: HSV-1 pUL28 (green), pUL15 (light blue), and pUL33 (light purple). Unmodelled aa residues, expected to comprise the unfolded region of each protein, were listed as follows: HSV-1 pUL28 (aa 268-305, 433-477, and 652-653), pUL15 (aa 178-194, 603-613, and 686-704), and pUL33 (aa 83-95). (B) The KSHV monomeric terminase complex was predicted by AlphaFold2. Each KSHV protein cartoon was colored as follows: KSHV ORF7 (green), ORF29 (light blue), and ORF67.5 (light purple). The KSHV terminase complex proteins were colored according to their HSV-1 homologs. (C) Based on the predicted KSHV terminase complex model, the main chain cartoon of the KSHV ORF67.5 regions substituted with alanine in the present study was visualized with rotation. The mutated regions (m1, m7, and m13), which are dispensable for the function of ORF67.5 in virus production, are highlighted in yellow. The mutated regions (m2-6 and m8-12) that are essential for the function of ORF67.5 in virus production are highlighted in magenta.

The interaction between ORF7 and ORF29 was promoted by ORF67.5, and the interaction between ORF67.5 and ORF7 was enhanced by ORF29 (Figs. 9A and B). In addition, we detected neither an ORF67.5-dependent increase in ORF7 and ORF29 expression, nor an ORF29-dependent increase in ORF67.5 and ORF7 expression (Figs. 9A and B). These data indicate that the increase in each protein-protein interaction is not due to an increase in expression nor stabilization of the two interacting proteins. Instead, the mechanism involves the enhancement or stabilization of the protein-protein interactions. As for the HSV-1 terminase complex components, UL28 (KSHV ORF7 homolog) directly interacts with UL15 (KSHV ORF29 homolog) and UL33 (KSHV ORF67.5 homolog) (22, 23). Similarly to KSHV ORF67.5, UL33 increases the interaction between UL28 and UL15. However, unlike KSHV ORF29, UL15 has no effect on the interaction between UL33 and UL28 (23). These results show that HSV-1 and KSHV homologs share some common features and differences. Specifically, the interaction between UL28 (ORF7) and UL15 (ORF29) is supported by UL33 (ORF67.5). On the other hand, in HSV-1, the interaction between UL33 and UL28 is not supported by UL15, but in KSHV, the interaction between ORF67.5 and ORF7 was increased in an ORF29 expression level-dependent manner. In addition, it has been reported that HSV-1 UL28 protects UL33 from proteasomal degradation (23), whereas our data showed that ORF67.5 was not stabilized by ORF7 (Fig. 8).

The ORF67.5 non-conserved region mutants (ORF67.5 m1, m7, and m13) retained the ability to complement the reduced virus production caused by ORF67.5 deletion and retained the ability to interact with ORF7 in the absence of ORF29 (Figs. 7 and 8). On the other hand, all the conserved region mutants of ORF67.5 (ORF67.5 m2-6 and m8-12) lost both the ability to complement virus production and the ability to interact with ORF7 in the absence of ORF29. However, among the conserved region mutants, ORF67.5 m9, m11, and m12 interacted with ORF7 in the presence of ORF29 (Fig. 10A). These findings suggest that the ability of ORF67.5 to interact with ORF7 in the presence of ORF29 is necessary, but not sufficient, for the contribution of ORF67.5 to virus production. Namely, the ability of ORF67.5 to interact with ORF7 in the absence of ORF29 may be sufficient for ORF67.5 to exert the ORF67.5-mediated function in virus production. Interestingly, it has been reported that a C-terminal region of HSV-1 UL33 is important for virus production but is not required for interaction with UL28 (47). This report prompted us to ask the following question: why were the ORF67.5 conserved region mutants (ORF67.5 m9, m11, and m12) defective in the ability to complement virus production despite interacting with ORF7 in the presence of ORF29? To answer this question, we propose the following three possibilities: (i) The ORF67.5 mutants (ORF67.5 m9, m11, and m12) can form a complex with ORF7 and ORF29, but the complex cannot adopt the proper conformation to function as a terminase; (ii) ORF67.5 has an unknown function in virus production that is absent in the mutants (ORF67.5 m9, m11, and m12) and (iii) The mutants (ORF67.5 m9, m11, and m12) are unable to interact with unknown elements important for terminase function other than ORF7 and ORF29. The answer to this question is unclear and requires future clarification. Moreover, a similar question needs to be addressed for HSV-1 UL33.

We have focused our studies on the components of the KSHV terminase complex and analyzed the functions of these proteins by constructing genetically modified viruses based on reverse genetics. Our previous studies have shown that ORF7 was important for KSHV terminase function (32, 33). Moreover, characterization of ORF67.5 and its contribution to terminase function was also accomplished to some extent in this study. The remaining important issue is to elucidate the role of ORF29 in terminase function. The C-terminal exon region of ORF29 has been reported to possess DNA sequence-independent nuclease activity (48). However, the function of ORF29 in virus replication is still unknown, and this functional analysis is currently underway as our next research project.

## Materials and methods

### Plasmids

The C-terminal 2×S-tagged ORF7 expression plasmid and the N-terminal 5×HA-tagged ORF67.5 expression plasmid have been described previously (32). The N-terminal 3xFLAG-tagged ORF67.5 expression plasmid was constructed using the insert extracted from a previously constructed N-terminal 2×S-tagged ORF67.5 expression plasmid digested with EcoRI and SalI (Takara Bio, Shiga, Japan) (32). The C-terminal 3xFLAG-tagged ORF29 expression plasmid was constructed with inserts obtained by PCR using the previously constructed N-terminal 3xFLAG-tagged ORF29 expression plasmid as a template (32). All ORF67.5 mutants were constructed by PCR using an N-terminal 3xFLAG-tagged ORF67.5 expression plasmid as a template. The KOD-Plus-Neo (TOYOBO, Osaka, Japan) was used for PCR, the DNA Ligation kit, Mighty Mix (Takara Bio) was used for ligation, and the pCI-neo mammalian expression vector (Promega, WI, USA) was used as the backbone vector. The primers used for plasmid construction are listed in Table 1. The sequences of the inserts were verified by Sanger sequencing.

**Table 1:**
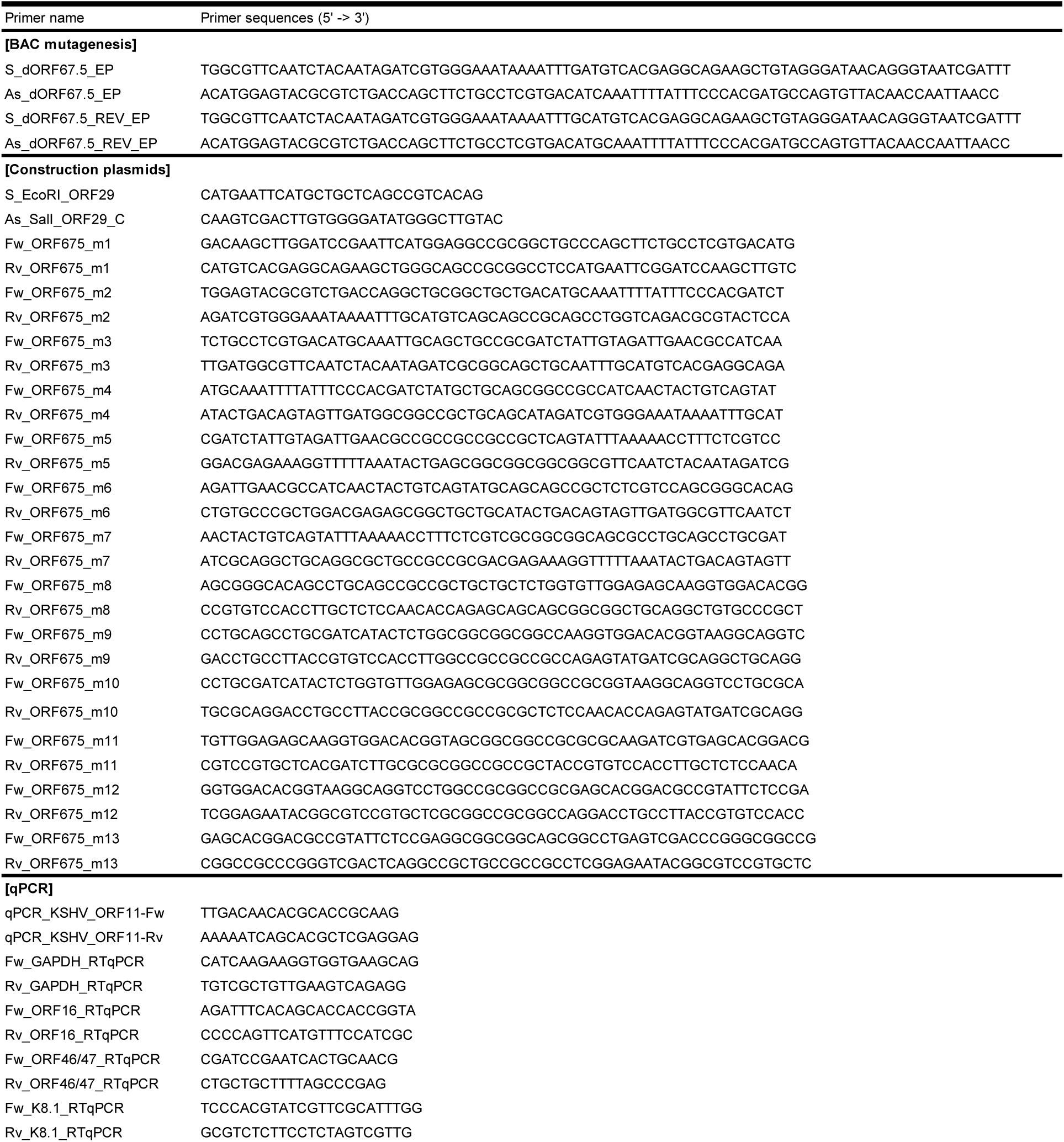
Primers for BAC mutagenesis, construction of plasmids, and qPCR

### Construction of ΔORF67.5-BAC16 and Revertant-BAC16

ΔORF67.5-BAC16 was generated from WT-BAC16 using a two-step markerless Red recombination system according to published protocols (39, 49, 50). ΔORF67.5-BAC16 contained a single cytosine deletion at position 114,586 of WT-BAC16 (GenBank: GQ994935). The primers used for this mutagenesis are shown in Table 1. The insertions and deletions of the kanamycin-resistance cassette in each mutant were analyzed by EcoRV digestion and agarose gel electrophoresis. The nucleotide sequence of the region surrounding the mutated site was determined by Sanger sequencing.

### Establishment of KSHV-BAC-harboring iSLK cells

In order to obtain efficient recombinant KSHV-producing cells, tetracycline (Tet)/ Dox-inducible (Tet-On) RTA/ORF50-expressing iSLK cells were used as virus-producing cells (38). The iSLK cells were cultured in Dulbecco’s modified Eagle’s medium (Nacalai Tesque Inc., Kyoto, Japan) supplemented with 10% fetal bovine serum, 1 μg/mL of puromycin (InvivoGen, CA, USA), and 0.25 mg/mL of G418 (Nacalai Tesque Inc.). Thirty-six μg of WT-BAC16 or its mutants (ΔORF67.5-BAC16 and Revertant-BAC16) were transfected into iSLK cells (1×10^6^ cells) by the calcium phosphate method. The transfected cells were selected under 1 mg/mL of hygromycin B (Wako, Osaka, Japan) to establish Dox-inducible recombinant KSHV-producing cell lines (WT-iSLK, ΔORF67.5-iSLK, and Revertant-iSLK).

### Characterization of mutant KSHVs

Each measurement was performed using previously described methods (33). Briefly, iSLK cells harboring WT or each KSHV mutant were treated with 8 μg/mL of Dox and 1.5 mM of SB for 72 h to induce lytic replication.

For measurement of viral gene expression, lytic-induced or uninduced cells (5×10^5^ cells on a 6-well plate) were harvested with 500 μL of RNAiso Plus (Takara Bio). Total RNA was extracted, and cDNA was synthesized from 160 ng of total RNA using the ReverTra Ace qPCR RT Master Mix (TOYOBO). RT-qPCR was performed using the synthesized cDNAs as template.

For quantification of intracellular viral DNA replication, iSLK cells (3.5×10^4^ cells on a 48-well plate) were induced or not induced and harvested. Viral genomic DNA was purified from the harvested cells using a QIAamp DNA Blood mini kit (Qiagen, CA, USA). The viral genome copy number was quantified by qPCR and normalized to the amount of total DNA.

For quantification of extracellular encapsidated viral DNA, iSLK cells (1.5×10^5^ cells on a 12-well plate) were induced, and the culture supernatants were harvested and centrifuged to remove debris. The supernatants were treated with DNase I (New England Biolabs, MA, USA), and encapsidated viral DNA was extracted from the supernatants using the QIAamp DNA Blood mini kit (Qiagen). The purified KSHV genomic DNA was quantified by qPCR.

For measurement of infectious virus, the supernatant transfer assay was performed. iSLK cells (1×10^6^ cells on a 6 cm dish) were induced, and the culture supernatants and cells were collected. The supernatants and cells were centrifuged, and the supernatant was mixed with trypsinized HEK293T cells (7.5×10^5^ cells). Polybrene (8 μg/mL) (Sigma-Aldrich, MO, USA) was added to the cell mixtures, which were then placed into well plates. The plates were centrifuged at 1,200*g* for 2 h at room temperature and incubated for 24 h at 37°C. GFP-positive cells were analyzed with a flow cytometer (FACSCalibur, Beckton Dickinson, CA, USA) using the CellQuest Pro software (Beckton Dickinson).

### qPCR

All qPCRs were performed using the THUNDERBIRD Next SYBR qPCR Mix (TOYOBO). In order to measure the KSHV genome copy number, qPCR was performed using the KSHV-encoded ORF11-specific primers shown in Table 1. Table 1 also lists the primers used to measure viral gene expression. The relative mRNA expression levels were determined by ΔΔCt methods and were normalized to GAPDH mRNA levels.

### Complementation assay

The complementation assay was performed as previously described (33). Briefly, iSLK cells were transfected with each plasmid using ScreenFect A-plus (Wako) and simultaneously cultured with medium containing 8 μg/mL of Dox and 1.5 mM of SB. After 72 h, virus production and infectivity were assessed according to the methods described in the section entitled “Characterization of mutant KSHVs”.

### Western blotting and antibodies

Western blotting was performed as previously described (51). Briefly, cells were washed with PBS, lysed with SDS sample buffer (containing 2% vol/vol 2-mercaptoethanol), sonicated for 10 s, reduced at 60°C for 10 min, and subjected to SDS-PAGE and Western blotting. When lytic-induced cells were used, the cells were treated with 8 μg/mL of Dox and 1.5 mM of SB for 72 h. Anti-KSHV ORF67.5 rabbit pAb was generated by GL Biochem, Shanghai, China, using the synthetic peptide Cys-KIVSTDAVFSEARAR (ORF67.5: aa 65-79) as the antigen. Anti-KSHV ORF67.5 rabbit pAb was purified from the immunized rabbit serum using antigen peptide affinity chromatography. The following primary antibodies were used: anti-K-bZIP mouse monoclonal antibody (mAb) (F33P1; Santa Cruz Biotechnology, TX, USA), anti-K8.1 A/B mouse mAb (4A4; Santa Cruz Biotechnology), anti-ORF21 rabbit pAb (previously produced in our laboratory) (52), anti-β-actin mouse mAb (AC-15; Santa Cruz Biotechnology), anti-FLAG mouse mAb (FLA-1; MBL, Nagoya, Japan), anti-S rabbit pAb (sc-802; Santa Cruz Biotechnology), and anti-HA mouse mAb (TANA2; MBL). Anti-mouse IgG-horseradish peroxidase (HRP) (NA931; Cytiva, Tokyo, Japan) and anti-rabbit IgG-HRP (#7074; Cell Signaling Technology, MA, USA) were used as the secondary antibodies.

### Pull-down assay

HEK293T cells (2×10^6^ cells on a 10 cm dish) were transfected with 12 μg of plasmid DNA and 36 μg of polyethylenimine hydrochloride/PEI MAX (Polysciences, Inc., PA, USA). The cells were then cultured for 20 h. Next, the cells were lysed with 1.5 mL of lysis buffer [50 mM Tris-HCl (pH 8.0), 120 mM NaCl, glycerol (1% vol/vol), Nonidet P-40 (0.2% vol/vol), and 1 mM dithiothreitol]. The cell extracts were incubated with 20 µL of S-protein-immobilized agarose beads for 1 h, and the beads were washed three times with lysis buffer. The washed beads were resuspended in 15 µL of SDS-PAGE sample buffer containing 2-mercaptoethanol (2% vol/vol). The precipitates were detected by Western blotting.

### Electron microscopy

In order to observe intracellular capsid formation, iSLK cells were cultured for 48 h in medium containing 8 μg/mL of Dox and 1.5 mM of SB. The cultured cells induced for virus production were washed with PBS and trypsinized for cell detachment. The detached cells were resuspended in medium, washed with PBS, and pelleted. The samples were then fixed in 2% glutaraldehyde in PBS (pH 7.2) for 2 h on ice, followed by incubation in 1% osmium tetroxide in PBS at 4°C for 1.5 h. The fixed samples were washed five times in PBS and dehydrated in graded ethanol. The samples were then embedded in an epoxy resin (Nisshin EM Co., Ltd., Tokyo, Japan) and processed by routine electron microscopy. Ultrathin sections were prepared using a Porter Blum ultramicrotome (Reichert-Nissei ULTRACUT-N; Leica, Wetzlar, Germany) and mounted on a copper grid (200 mesh) supported by a carbon-coated collodion film. The ultrathin sections were double stained with uranyl acetate (4% wt/vol) and lead citrate for 10 min and 3 min, respectively. After each staining procedure, the sections were washed three times in distilled water. All sections were observed under a Hitachi H-7800 transmission electron microscope.

### Southern blotting

Southern blotting was performed as previously described (33, 37). Briefly, iSLK cells (2×10^6^ cells on a 10 cm dish) were treated with 8 μg/mL of Dox and 1.5 mM of SB for 72 h. Next, the cells were washed with PBS and suspended in TNE buffer [10 mM Tris-HCl (pH 7.5), 100 mM NaCl, and 1 mM EDTA], followed by an overnight incubation at 60°C with SDS (0.3% wt/vol) and proteinase K (100 μg/mL). The cell extracts were then incubated with RNase (100 μg/mL) at room temperature for 30 min. DNA was isolated from the cell extracts by phenol-chloroform extraction and ethanol precipitation. The pellets were then resuspended in 1×TE buffer. The DNA samples (10 μg each) were digested with EcoRI (Takara Bio) and SalI (TOYOBO) overnight, followed by electrophoresis on a 0.7% agarose 1×TBE gel. The gel was denatured in alkaline denaturing buffer (0.5 N NaOH and 1.5 M NaCl) for 45 min and transferred to a Hybond-N+ membrane (Cytiva) by capillary action overnight in the presence of alkaline transfer buffer (0.4 N NaOH and 1 M NaCl). The transferred DNA was cross-linked, and the membrane was treated with the DIG-High Prime DNA Labeling and Detection Starter kit II (Roche, Basel, Switzerland) according to the manufacturer’s protocol. The membrane was hybridized overnight at 42°C in DIG Easy Hyb buffer with a digoxigenin (DIG)-labeled TR probe. Next, the membrane was washed twice with 2×SSC, 0.1% SDS at room temperature followed by two additional washes with 0.5×SSC, 0.1% SDS at 68°C. The membrane was blocked, incubated with anti-DIG antibody, and detected by CSPD. The blot was then exposed to X-ray film (Fuji film, Tokyo, Japan).

### Protein structure prediction

The structure of the monomeric HSV-1 terminase complex (PDB; 6M5R) (22), determined by cryo-EM, was visualized with the molecular visualization open-source software program PyMOL (Ver 2.5.0). The structure of the monomeric KSHV terminase complex, determined by the deep learning algorithm AlphaFold 2.2.0 (https://github.com/deepmind/alphafold) in the local environment (53), was also visualized with PyMOL.

### Statistics

The statistical significance was determined by one-way ANOVA followed by Dunnett’s test and was evaluated using GraphPad Prism 7 software (GraphPad Software, CA, USA).

## Acknowledgements

The KSHV BAC clone, BAC16, was a kind gift from Jae U. Jung (Cleveland Clinic Lerner Research Institute, USA). We thank Gregory A. Smith (Northwestern University, USA) for the Escherichia coli strain GS1783, Yoshihiko Fujioka (Osaka Medical and Pharmaceutical University, Japan) for electron microscopy analysis, Rimiko Okabe (Fujimuro Lab. member) for assistance with data collection, and Nikolaus Osterrieder (Cornell University, USA) for the plasmid pEP-KanS. Y.I. was supported by the Nagai Memorial Research Scholarship from the Pharmaceutical Society of Japan and the JSPS Research Fellowship for Young Scientists. This work was partially supported by grants from the JSPS Grant-in-Aid for Scientific Research (18K06642; M.F. and JP22J23607; Y.I.).

## References

1) Chang Y, Cesarman E, Pessin MS, Lee F, Culpepper J, Knowles DM, Moore PS. Identification of herpesvirus-like DNA sequences in AIDS-associated Kaposi’s sarcoma. Science. 1994 Dec 16;266(5192):1865–9. doi: 10.1126/science.7997879. PMID: 7997879.

2) Damania B, Cesarman E. 2022. Kaposi’s Sarcoma Herpesvirus, p 513–572. In Howley PM, Knipe DM. (ed), Fields virology, 7th ed. Wolters Kluwer, Philadelphia, PA.

3) Russo JJ, Bohenzky RA, Chien MC, Chen J, Yan M, Maddalena D, Parry JP, Peruzzi D, Edelman IS, Chang Y, Moore PS. Nucleotide sequence of the Kaposi sarcoma-associated herpesvirus (HHV8). Proc Natl Acad Sci U S A. 1996 Dec 10;93(25):14862–7. doi: 10.1073/pnas.93.25.14862. PMID: 8962146; PMCID: PMC26227.

4) Nador RG, Cesarman E, Chadburn A, Dawson DB, Ansari MQ, Sald J, Knowles DM. Primary effusion lymphoma: a distinct clinicopathologic entity associated with the Kaposi’s sarcoma-associated herpes virus. Blood. 1996 Jul 15;88(2):645–56. PMID: 8695812.

5) Soulier J, Grollet L, Oksenhendler E, Cacoub P, Cazals-Hatem D, Babinet P, d’Agay MF, Clauvel JP, Raphael M, Degos L, et al. Kaposi’s sarcoma-associated herpesvirus-like DNA sequences in multicentric Castleman’s disease. Blood. 1995 Aug 15;86(4):1276–80. PMID: 7632932.

6) Uldrick TS, Wang V, O’Mahony D, Aleman K, Wyvill KM, Marshall V, Steinberg SM, Pittaluga S, Maric I, Whitby D, Tosato G, Little RF, Yarchoan R. An interleukin-6-related systemic inflammatory syndrome in patients co-infected with Kaposi sarcoma-associated herpesvirus and HIV but without Multicentric Castleman disease. Clin Infect Dis. 2010 Aug 1;51(3):350–8. doi: 10.1086/654798. PMID: 20583924; PMCID: PMC2946207.

7) Sun R, Lin SF, Gradoville L, Yuan Y, Zhu F, Miller G. A viral gene that activates lytic cycle expression of Kaposi’s sarcoma-associated herpesvirus. Proc Natl Acad Sci U S A. 1998 Sep 1;95(18):10866–71. doi: 10.1073/pnas.95.18.10866. PMID: 9724796; PMCID: PMC27987.

8) Brown JC, Newcomb WW. Herpesvirus capsid assembly: insights from structural analysis. Curr Opin Virol. 2011 Aug;1(2):142–9. doi: 10.1016/j.coviro.2011.06.003. PMID: 21927635; PMCID: PMC3171831.

9) Heming JD, Conway JF, Homa FL. Herpesvirus Capsid Assembly and DNA Packaging. Adv Anat Embryol Cell Biol. 2017;223:119–142. doi: 10.1007/978-3-319-53168-7_6. PMID: 28528442; PMCID: PMC5548147.

10) Newcomb WW, Homa FL, Thomsen DR, Booy FP, Trus BL, Steven AC, Spencer JV, Brown JC. Assembly of the herpes simplex virus capsid: characterization of intermediates observed during cell-free capsid formation. J Mol Biol. 1996 Nov 1;263(3):432–46. doi: 10.1006/jmbi.1996.0587. PMID: 8918599.

11) Heymann JB, Cheng N, Newcomb WW, Trus BL, Brown JC, Steven AC. Dynamics of herpes simplex virus capsid maturation visualized by time-lapse cryo-electron microscopy. Nat Struct Biol. 2003 May;10(5):334–41. doi: 10.1038/nsb922. PMID: 12704429.

12) Aksyuk AA, Newcomb WW, Cheng N, Winkler DC, Fontana J, Heymann JB, Steven AC. Subassemblies and asymmetry in assembly of herpes simplex virus procapsid. mBio. 2015 Oct 6;6(5):e01525–15. doi: 10.1128/mBio.01525-15. PMID: 26443463; PMCID: PMC4611051.

13) Gibson W, Roizman B. Proteins specified by herpes simplex virus. 8. Characterization and composition of multiple capsid forms of subtypes 1 and 2. J Virol. 1972 Nov;10(5):1044–52. doi: 10.1128/JVI.10.5.1044-1052.1972. PMID: 4344252; PMCID: PMC356576.

14) Gao M, Matusick-Kumar L, Hurlburt W, DiTusa SF, Newcomb WW, Brown JC, McCann PJ 3rd, Deckman I, Colonno RJ. The protease of herpes simplex virus type 1 is essential for functional capsid formation and viral growth. J Virol. 1994 Jun;68(6):3702–12. doi: 10.1128/JVI.68.6.3702-3712.1994. PMID: 8189508; PMCID: PMC236875.

15) Newcomb WW, Trus BL, Cheng N, Steven AC, Sheaffer AK, Tenney DJ, Weller SK, Brown JC. Isolation of herpes simplex virus procapsids from cells infected with a protease-deficient mutant virus. J Virol. 2000 Feb;74(4):1663–73. doi: 10.1128/jvi.74.4.1663-1673.2000. PMID: 10644336; PMCID: PMC111641.

16) Schrag JD, Prasad BV, Rixon FJ, Chiu W. Three-dimensional structure of the HSV1 nucleocapsid. Cell. 1989 Feb 24;56(4):651–60. doi: 10.1016/0092-8674(89)90587-4. PMID: 2537151.

17) Booy FP, Newcomb WW, Trus BL, Brown JC, Baker TS, Steven AC. Liquid-crystalline, phage-like packing of encapsidated DNA in herpes simplex virus. Cell. 1991 Mar 8;64(5):1007–15. doi: 10.1016/0092-8674(91)90324-r. PMID: 1848156; PMCID: PMC4140082.

18) Krug LT, Pellett PE. 2022. The Family Herpesviridae: A Brief Introduction, p 212–234. In Howley PM, Knipe DM. (ed), Fields virology, 7th ed. Wolters Kluwer, Philadelphia, PA.

19) Russo JJ, Bohenzky RA, Chien MC, Chen J, Yan M, Maddalena D, Parry JP, Peruzzi D, Edelman IS, Chang Y, Moore PS. Nucleotide sequence of the Kaposi sarcoma-associated herpesvirus (HHV8). Proc Natl Acad Sci U S A. 1996 Dec 10;93(25):14862–7. doi: 10.1073/pnas.93.25.14862. PMID: 8962146; PMCID: PMC26227.

20) Beard PM, Taus NS, Baines JD. DNA cleavage and packaging proteins encoded by genes U(L)28, U(L)15, and U(L)33 of herpes simplex virus type 1 form a complex in infected cells. J Virol. 2002 May;76(10):4785–91. doi: 10.1128/jvi.76.10.4785-4791.2002. PMID: 11967295; PMCID: PMC136146.

21) Heming JD, Huffman JB, Jones LM, Homa FL. Isolation and characterization of the herpes simplex virus 1 terminase complex. J Virol. 2014 Jan;88(1):225–36. doi: 10.1128/JVI.02632-13. Epub 2013 Oct 23. PMID: 24155374; PMCID: PMC3911699.

22) Yang Y, Yang P, Wang N, Chen Z, Su D, Zhou ZH, Rao Z, Wang X. Architecture of the herpesvirus genome-packaging complex and implications for DNA translocation. Protein Cell. 2020 May;11(5):339–351. doi: 10.1007/s13238-020-00710-0. Epub 2020 Apr 23. PMID: 32328903; PMCID: PMC7196598.

23) Yang K, Baines JD. The putative terminase subunit of herpes simplex virus 1 encoded by UL28 is necessary and sufficient to mediate interaction between pUL15 and pUL33. J Virol. 2006 Jun;80(12):5733–9. doi: 10.1128/JVI.00125-06. PMID: 16731912; PMCID: PMC1472570.

24) Poon AP, Roizman B. Characterization of a temperature-sensitive mutant of the UL15 open reading frame of herpes simplex virus 1. J Virol. 1993 Aug;67(8):4497–503. doi: 10.1128/JVI.67.8.4497-4503.1993. PMID: 8331721; PMCID: PMC237833.

25) Baines JD, Poon AP, Rovnak J, Roizman B. The herpes simplex virus 1 UL15 gene encodes two proteins and is required for cleavage of genomic viral DNA. J Virol. 1994 Dec;68(12):8118–24. doi: 10.1128/JVI.68.12.8118-8124.1994. PMID: 7966602; PMCID: PMC237276.

26) Addison C, Rixon FJ, Preston VG. Herpes simplex virus type 1 UL28 gene product is important for the formation of mature capsids. J Gen Virol. 1990 Oct;71 (Pt 10):2377–84. doi: 10.1099/0022-1317-71-10-2377. PMID: 2172450.

27) Tengelsen LA, Pederson NE, Shaver PR, Wathen MW, Homa FL. Herpes simplex virus type 1 DNA cleavage and encapsidation require the product of the UL28 gene: isolation and characterization of two UL28 deletion mutants. J Virol. 1993 Jun;67(6):3470–80. doi: 10.1128/JVI.67.6.3470-3480.1993. PMID: 8388510; PMCID: PMC237693.

28) al-Kobaisi MF, Rixon FJ, McDougall I, Preston VG. The herpes simplex virus UL33 gene product is required for the assembly of full capsids. Virology. 1991 Jan;180(1):380–8. doi: 10.1016/0042-6822(91)90043-b. PMID: 1845831.

29) Patel AH, Rixon FJ, Cunningham C, Davison AJ. Isolation and characterization of herpes simplex virus type 1 mutants defective in the UL6 gene. Virology. 1996 Mar 1;217(1):111–23. doi: 10.1006/viro.1996.0098. PMID: 8599195.

30) Goldner T, Hewlett G, Ettischer N, Ruebsamen-Schaeff H, Zimmermann H, Lischka P. The novel anticytomegalovirus compound AIC246 (Letermovir) inhibits human cytomegalovirus replication through a specific antiviral mechanism that involves the viral terminase. J Virol. 2011 Oct;85(20):10884–93. doi: 10.1128/JVI.05265-11. Epub 2011 Jul 13. PMID: 21752907; PMCID: PMC3187482.

31) Marty FM, Ljungman P, Chemaly RF, Maertens J, Dadwal SS, Duarte RF, Haider S, Ullmann AJ, Katayama Y, Brown J, Mullane KM, Boeckh M, Blumberg EA, Einsele H, Snydman DR, Kanda Y, DiNubile MJ, Teal VL, Wan H, Murata Y, Kartsonis NA, Leavitt RY, Badshah C. Letermovir Prophylaxis for Cytomegalovirus in Hematopoietic-Cell Transplantation. N Engl J Med. 2017 Dec 21;377(25):2433–2444. doi: 10.1056/NEJMoa1706640. Epub 2017 Dec 6. PMID: 29211658.

32) Iwaisako Y, Watanabe T, Hanajiri M, Sekine Y, Fujimuro M. Kaposi’s Sarcoma-Associated Herpesvirus ORF7 Is Essential for Virus Production. Microorganisms. 2021 May 28;9(6):1169. doi: 10.3390/microorganisms9061169. PMID: 34071710; PMCID: PMC8228664.

33) Iwaisako Y, Watanabe T, Futo M, Okabe R, Sekine Y, Suzuki Y, Nakano T, Fujimuro M. The Contribution of Kaposi’s Sarcoma-Associated Herpesvirus ORF7 and Its Zinc-Finger Motif to Viral Genome Cleavage and Capsid Formation. J Virol. 2022 Sep 28;96(18):e0068422. doi: 10.1128/jvi.00684-22. Epub 2022 Sep 8. PMID: 36073924; PMCID: PMC9517700.

34) Vizoso Pinto MG, Pothineni VR, Haase R, Woidy M, Lotz-Havla AS, Gersting SW, Muntau AC, Haas J, Sommer M, Arvin AM, Baiker A. Varicella zoster virus ORF25 gene product: an essential hub protein linking encapsidation proteins and the nuclear egress complex. J Proteome Res. 2011 Dec 2;10(12):5374–82. doi: 10.1021/pr200628s. Epub 2011 Oct 26. PMID: 21988664; PMCID: PMC3230707.

35) Borst EM, Kleine-Albers J, Gabaev I, Babic M, Wagner K, Binz A, Degenhardt I, Kalesse M, Jonjic S, Bauerfeind R, Messerle M. The human cytomegalovirus UL51 protein is essential for viral genome cleavage-packaging and interacts with the terminase subunits pUL56 and pUL89. J Virol. 2013 Feb;87(3):1720–32. doi: 10.1128/JVI.01955-12. Epub 2012 Nov 21. PMID: 23175377; PMCID: PMC3554196.

36) Arias C, Weisburd B, Stern-Ginossar N, Mercier A, Madrid AS, Bellare P, Holdorf M, Weissman JS, Ganem D. KSHV 2.0: a comprehensive annotation of the Kaposi’s sarcoma-associated herpesvirus genome using next-generation sequencing reveals novel genomic and functional features. PLoS Pathog. 2014 Jan;10(1):e1003847. doi: 10.1371/journal.ppat.1003847. Epub 2014 Jan 16. PMID: 24453964; PMCID: PMC3894221.

37) Gardner MR, Glaunsinger BA. Kaposi’s Sarcoma-Associated Herpesvirus ORF68 Is a DNA Binding Protein Required for Viral Genome Cleavage and Packaging. J Virol. 2018 Jul 31;92(16):e00840–18. doi: 10.1128/JVI.00840-18. PMID: 29875246; PMCID: PMC6069193.

38) Myoung J, Ganem D. Generation of a doxycycline-inducible KSHV producer cell line of endothelial origin: maintenance of tight latency with efficient reactivation upon induction. J Virol Methods. 2011 Jun;174(1-2):12–21. doi: 10.1016/j.jviromet.2011.03.012. Epub 2011 Mar 17. PMID: 21419799; PMCID: PMC3095772.

39) Brulois KF, Chang H, Lee AS, Ensser A, Wong LY, Toth Z, Lee SH, Lee HR, Myoung J, Ganem D, Oh TK, Kim JF, Gao SJ, Jung JU. Construction and manipulation of a new Kaposi’s sarcoma-associated herpesvirus bacterial artificial chromosome clone. J Virol. 2012 Sep;86(18):9708–20. doi: 10.1128/JVI.01019-12. Epub 2012 Jun 27. PMID: 22740391; PMCID: PMC3446615.

40) Deng B, O’Connor CM, Kedes DH, Zhou ZH. Cryo-electron tomography of Kaposi’s sarcoma-associated herpesvirus capsids reveals dynamic scaffolding structures essential to capsid assembly and maturation. J Struct Biol. 2008 Mar;161(3):419–27. doi: 10.1016/j.jsb.2007.10.016. Epub 2007 Nov 17. PMID: 18164626; PMCID: PMC2692512.

41) Arvin AM, Abendroth A. 2022. Varicella-zoster virus, p 445–488. In Howley PM, Knipe DM. (ed), Fields virology, 7th ed. Wolters Kluwer, Philadelphia, PA.

42) Dai X, Gong D, Lim H, Jih J, Wu TT, Sun R, Zhou ZH. Structure and mutagenesis reveal essential capsid protein interactions for KSHV replication. Nature. 2018 Jan 25;553(7689):521–525. doi: 10.1038/nature25438. Epub 2018 Jan 17. PMID: 29342139; PMCID: PMC6039102.

43) Nealon K, Newcomb WW, Pray TR, Craik CS, Brown JC, Kedes DH. Lytic replication of Kaposi’s sarcoma-associated herpesvirus results in the formation of multiple capsid species: isolation and molecular characterization of A, B, and C capsids from a gammaherpesvirus. J Virol. 2001 Mar;75(6):2866–78. doi: 10.1128/JVI.75.6.2866-2878.2001. PMID: 11222712; PMCID: PMC115913.

44) Visalli MA, House BL, Lahrman FJ, Visalli RJ. Intermolecular Complementation between Two Varicella-Zoster Virus pORF30 Terminase Domains Essential for DNA Encapsidation. J Virol. 2015 Oct;89(19):10010–22. doi: 10.1128/JVI.01313-15. Epub 2015 Jul 22. PMID: 26202238; PMCID: PMC4577918.

45) Tsurumi S, Watanabe T, Iwaisako Y, Suzuki Y, Nakano T, Fujimuro M. Kaposi’s sarcoma-associated herpesvirus ORF17 plays a key role in capsid maturation. Virology. 2021 Jun;558:76–85. doi: 10.1016/j.virol.2021.02.009. Epub 2021 Feb 25. PMID: 33735753.

46) Cardone G, Moyer AL, Cheng N, Thompson CD, Dvoretzky I, Lowy DR, Schiller JT, Steven AC, Buck CB, Trus BL. Maturation of the human papillomavirus 16 capsid. mBio. 2014 Aug 5;5(4):e01104–14. doi: 10.1128/mBio.01104-14. PMID: 25096873; PMCID: PMC4128349.

47) Beilstein F, Higgs MR, Stow ND. Mutational analysis of the herpes simplex virus type 1 DNA packaging protein UL33. J Virol. 2009 Sep;83(17):8938–45. doi: 10.1128/JVI.01048-09. Epub 2009 Jun 24. PMID: 19553324; PMCID: PMC2738152.

48) Miller JT, Zhao H, Masaoka T, Varnado B, Cornejo Castro EM, Marshall VA, Kouhestani K, Lynn AY, Aron KE, Xia A, Beutler JA, Hirsch DR, Tang L, Whitby D, Murelli RP, Le Grice SFJ. Sensitivity of the C-Terminal Nuclease Domain of Kaposi’s Sarcoma-Associated Herpesvirus ORF29 to Two Classes of Active-Site Ligands. Antimicrob Agents Chemother. 2018 Sep 24;62(10):e00233–18. doi: 10.1128/AAC.00233-18. PMID: 30061278; PMCID: PMC6153795.

49) Tischer BK, Smith GA, Osterrieder N. En passant mutagenesis: a two step markerless red recombination system. Methods Mol Biol. 2010;634:421–30. doi: 10.1007/978-1-60761-652-8_30. PMID: 20677001.

50) Watanabe T, Nishimura M, Izumi T, Kuriyama K, Iwaisako Y, Hosokawa K, Takaori-Kondo A, Fujimuro M. Kaposi’s Sarcoma-Associated Herpesvirus ORF66 Is Essential for Late Gene Expression and Virus Production via Interaction with ORF34. J Virol. 2020 Jan 6;94(2):e01300–19. doi: 10.1128/JVI.01300-19. PMID: 31694948; PMCID: PMC6955251.

51) Fujimuro M, Wu FY, ApRhys C, Kajumbula H, Young DB, Hayward GS, Hayward SD. A novel viral mechanism for dysregulation of beta-catenin in Kaposi’s sarcoma-associated herpesvirus latency. Nat Med. 2003 Mar;9(3):300–6. doi: 10.1038/nm829. Epub 2003 Feb 18. PMID: 12592400.

52) Yamaguchi T, Watanabe T, Iwaisako Y, Fujimuro M. Kaposi’s Sarcoma-Associated Herpesvirus ORF21 Enhances the Phosphorylation of MEK and the Infectivity of Progeny Virus. Int J Mol Sci. 2023 Jan 8;24(2):1238. doi: 10.3390/ijms24021238. PMID: 36674756; PMCID: PMC9867424.

53) Jumper J, Evans R, Pritzel A, Green T, Figurnov M, Ronneberger O, Tunyasuvunakool K, Bates R, Žídek A, Potapenko A, Bridgland A, Meyer C, Kohl SAA, Ballard AJ, Cowie A, Romera-Paredes B, Nikolov S, Jain R, Adler J, Back T, Petersen S, Reiman D, Clancy E, Zielinski M, Steinegger M, Pacholska M, Berghammer T, Bodenstein S, Silver D, Vinyals O, Senior AW, Kavukcuoglu K, Kohli P, Hassabis D. Highly accurate protein structure prediction with AlphaFold. Nature. 2021 Aug;596(7873):583–589. doi: 10.1038/s41586-021-03819-2. Epub 2021 Jul 15. PMID: 34265844; PMCID: PMC8371605.

